# Integrating Proteomes for Lung Tissue and Lavage Reveals Pathways that Link Responses in Allergen-Challenged Mice

**DOI:** 10.1101/2020.01.14.905737

**Authors:** Thomas H Mahood, Christopher D Pascoe, Tobias K Karakach, Aruni Jha, Sujata Basu, Peyman Ezzati, Victor Spicer, Neeloffer Mookherjee, Andrew J Halayko

## Abstract

To capture interplay between biological pathways we analyzed the proteome from matched lung tissue and bronchoalveolar lavage fluid (BALF) of individual allergen-naïve and house dust mite (HDM)-challenged BALB/c mice, a model of allergic asthma. Unbiased label-free LC-MS/MS analysis quantified 2,675 proteins from tissue and BALF of allergen-naïve and HDM-exposed mice. In comparing the four datasets we found significantly greater diversity in proteins between lung tissue and BALF than in the changes induced by HDM challenge. The biological pathways enriched after allergen exposure were compartment-dependent. Lung tissue featured innate immune responses and oxidative stress, while BALF most strongly revealed changes in metabolism. We combined lung tissue and BALF proteomes, which principally highlighted oxidation reduction (redox) pathways, a finding influenced chiefly by the lung tissue dataset. Integrating lung and BALF proteomes also uncovered new proteins and biological pathways that may mediate lung tissue and BALF interactions after allergen challenge, for example, B Cell Receptor signaling. We demonstrate that enhanced insight is fostered when different biological compartments from the lung are investigated in parallel. Integration of proteomes from lung tissue and BALF compartments reveals new information about protein networks in the response to environmental challenge and interaction between intracellular and extracellular process.

## Introduction

The biological underpinnings of chronic inflammation in asthma involve a complex molecular interplay between structural cells, recruited inflammatory cells, and the extracellular mediators that they release. Understanding this interplay is important to understand pathobiology and support pre-clinical research to develop new therapies. Mice exposed to allergen are a mainstay for asthma research, and a common approach is to challenge animals with inhaled house dust mite (HDM). HDM is an aeroallergen that is clinically relevant, and is a complex stimulus with multiple immunogens and stressors, including fungal spores, bacterial endotoxins, lipid-binding proteins and proteases^1, 2^. Repeated HDM challenge induces pathophysiological symptoms that are hallmarks of human asthma^3^. However, the scope and complexity of the interplay between lung cells, autocrine and paracrine pathways, and the integrated signaling networks associated with the pathobiology that determines the efficacy of pre-clinical therapeutics, has not been refined.

To understand complex disease mechanisms, a number of omics technologies have been employed, including proteomics. To the best of our knowledge, most studies examine the proteome of individual biological compartments, including airway spaces - collected as bronchoalveolar lavage fluid (BALF) or sputum - and lung tissue from asthmatic patients and murine models of the disease^4–6^. These studies have been important for endotyping patients and animal models, identifying biomarkers of disease, and providing direction for new therapeutic strategies. However, analysis of these biological compartments in isolation can only partially identify the molecular networks and biomarkers that are critical for disease expression in a complex system like the lung. Nonetheless, understanding the molecular systems that determine the interaction between secreted proteins in the lung and the pathways that are regulated in the resident cells of the lung has not been fully established in conventional proteomics studies.

To address this knowledge gap, we used unbiased proteomics and unsupervised exploratory data analysis to establish a molecular signature in matched lung tissue and BALF from individual mice. We then combined these to create integrated proteomes that allow unique signals from each sample type to be identified, and the nature of the interactions between biological compartments to be deciphered in describing the response to HDM challenge. We found that the proteomes of lung tissue and of BALF are distinct and that inhaled HDM challenge induces unique alterations in each compartment. Moreover, by integrating proteomes we uncovered novel signaling networks that were not evident from individual proteome dataset of lung tissue or BALF.

## Results

### Lung function & differential cell count analysis

HDM challenge of adult female BALB/c mice resulted in pathophysiological features that are consistent with human asthma. This included significant increases in airway resistance, tissue elastance and tissue resistance in methacholine challenge (Supplemental Figure S1A). Differential counting of BALF immune cells confirmed that HDM challenge triggered significant accumulation of eosinophils and neutrophils that is also consistent with prior studies^3^ and mimics human asthma (Supplemental Figure S1B).

### Lung tissue and BALF proteome disparities

In total, our analysis of lung tissue and BALF samples in all mice, yielded 2,675 uniquely identifiable proteins (Protein IDs). On a per mouse basis, we obtained 1,594 ± 154 protein IDs (mean ± SD) from lung tissue and 641 ± 207 protein IDs from BALF samples (Supplemental Figure S2A). To confirm that secreted proteins were enriched in the BALF samples, we characterized protein IDs using the Uniprot keyword, “secreted”, which annotated >75% of the proteins in the BALF. In contrast, ~only 25% of the protein ID’s were characterized as “secreted” in the tissue lysate. To confirm that membrane-associated proteins are enriched in lung tissue samples, we classified proteins using the Uniprot annotated keyword “transmembrane”. This identified 69% of proteins in lung tissue, and only 23% of BALF proteins.

To investigate if the proteome of lung tissue and BALF from allergen-naïve and HDM-exposed mice are distinct, we performed Principal Component Analysis (PCA) on the top 500 most variable proteins across all samples. This cut-off was chosen arbitrarily to capture the high intensity (and therefore highly variable) signals from our dataset. Our comparison matrix of all samples from allergen-naïve and HDM-challenged mice using the top two principal components (PC) accounted for ~80% of variability between samples (Figure 1A). Lung tissue and BALF samples are separated along the 1^st^ PC (x-axis), which accounts for approximately 59% of the data variance. Lung tissue and BALF samples from naïve and HDM-challenged mice were also distinct, with the 2^nd^ PC accounting for about 20% of data variance, with HDM effects seemingly much greater in BALF than in lung tissue (Figure 1A, expanded panel).

**Figure 1:**
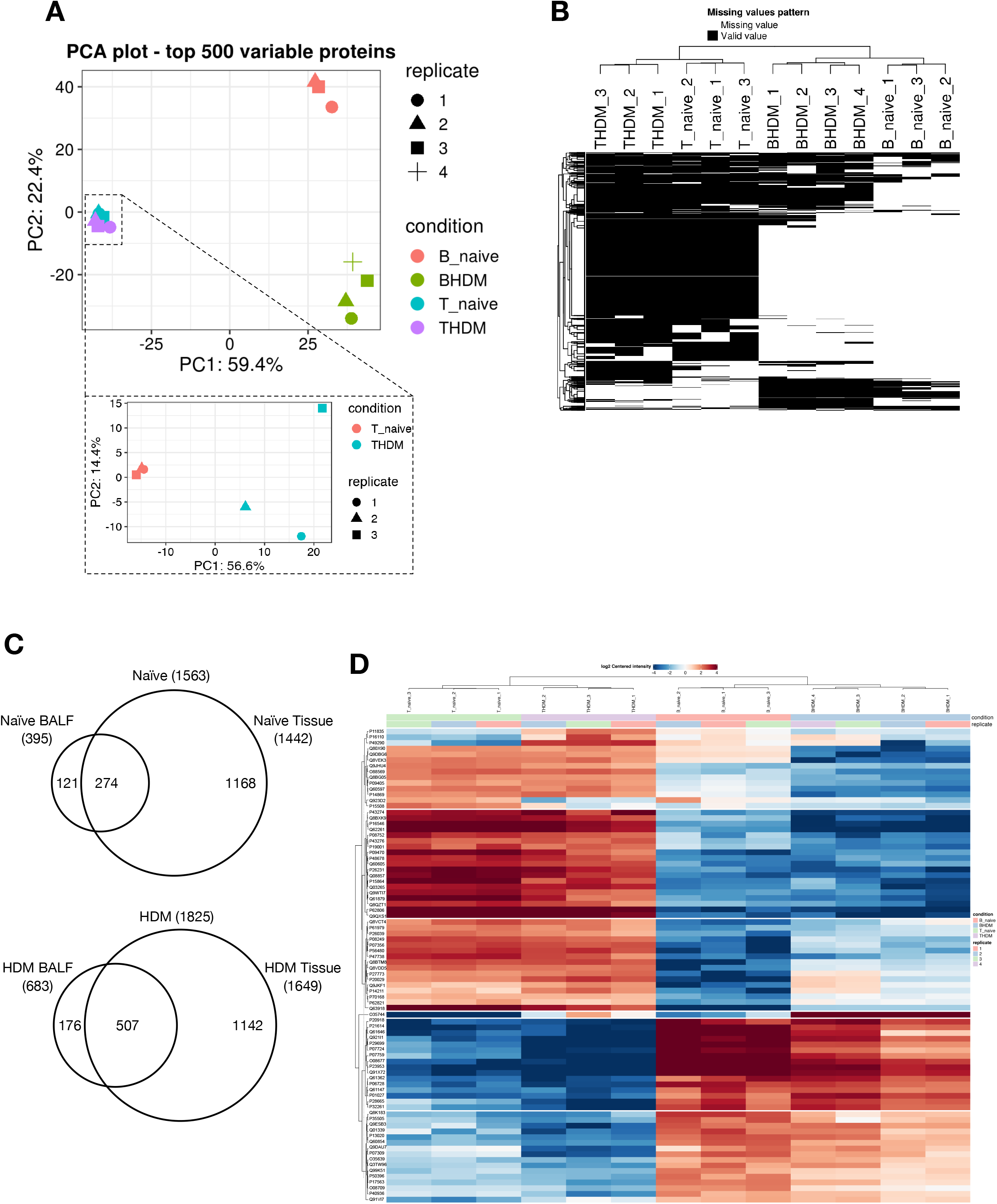
Characterization of the distinct proteomic signatures of the lung tissue and BALF. **A)** PCA of the top 500 most variable proteins identifies that BALF and tissue datasets have greater variation (59.4%) than the datasets comparing allergen-naïve and HDM challenged samples (22.4%). Note that the variability between the highly divergent tissue and BALF datasets masks the differences shown between allergen-naïve and HDM-exposed lung tissue as described in the figure inset (PCA of the top 500 most variable proteins). **B)** The differences between BALF and tissue samples is driven by missing proteins in BALF compartment as shown by K-means clustering heatmap of detectable proteins. **C)** The distribution of Uniprot ID’s across lung tissue and BALF in both naïve and HDM exposed mice. **D)** K-means clustering heatmap showing protein abundance differences between lung tissue and BALF from allergen naive and HDM mice. K-means clustering identified 6 clusters (numbered on the left hand side). Red – high abundance, Blue – low abundance. Abbreviations used: House Dust Mite (HDM), Bronchial Alveolar Lavage Fluid (BALF), Principal Component Analysis (PCA).

We next examined the distribution of proteins across all samples. A clustering heatmap (k-means) shows that BALF and tissue clustered separately, as did naïve and HDM-challenged for tissue and BALF (Figure 1B). We also observed that the differences in lung tissue and BALF samples in allergen-naïve and HDM-challenged mice were associated with the unique protein IDs in individual datasets. Approximately 72.7% of all protein IDs were unique to either lung tissue or BALF samples.

We next examined the distribution of Uniprot IDs in lung tissue and BALF, and the effects of HDM-challenge (Figure 1C). Venn diagrams show that HDM exposure increased the absolute number of proteins in lung tissue and BALF by 114% and 173%, respectively. The fraction of protein IDs common to BALF and lung tissue was 18% in allergen-naïve mice, and this increased to 28% after HDM exposure. HDM exposure reduced the fraction of unique lung tissue Uniprot IDs from 75% to 63%. The proportion of proteins that were unique to BALF was 8-10%; this was not changed by HDM-challenge.

We next examined patterns of protein abundance of the specific subset that were common to BALF and lung tissue from allergen-naïve and HDM-challenged mice (Figure 1D). The dominating feature of the heat map is that it discerns large clusters of proteins that are relatively more prominent in BALF or lung tissue, independent of the effects of HDM challenge. Thus, for these proteins that are common to lung and BALF, k-means clustering is consistent with PC analysis of all proteins (Figure 1A) which showed that data variability was chiefly the result of differences between lung tissue and BALF. Though effects of HDM challenge within lung tissue or BALF are evident in this common protein subset, our analysis highlights the disparities between lung and BALF that likely affect the biological manifestation of HDM challenge.

### Independent validation of proteins selected from the proteomic analysis

To validate our proteomic data, immunoblotting was performed on the same samples for three proteins (Figure 2). ARG1 (Arginase-1), CLCA1 (Calcium-activated chloride channel regulator 1) and FDPS (Farnesyl pyrophosphate synthase) were selected based upon their enrichment by HDM exposure and the availability of reliable immunoblotting grade antibodies. Immunoblotting confirmed our proteomic data for ARG1 and CLCA1, as they were undetectable in allergen-naïve samples from lung tissue and BALF, but prominent bands were evident for samples from allergen-challenged mice (Figures 2A and 2B). Immunoblotting analysis confirmed presence of FDPS in all samples, with a significant enrichment of 270% (p.adj ≤ 0.05) in the lung tissue after HDM-challenge (Figures 2A and 2B). This is consistent with proteomic data for FDPS, which was detected in only one BALF and lung tissue sample from an allergen-naïve mouse, while it was present and increased in all samples from HDM-challenged mice.

**Figure 2:**
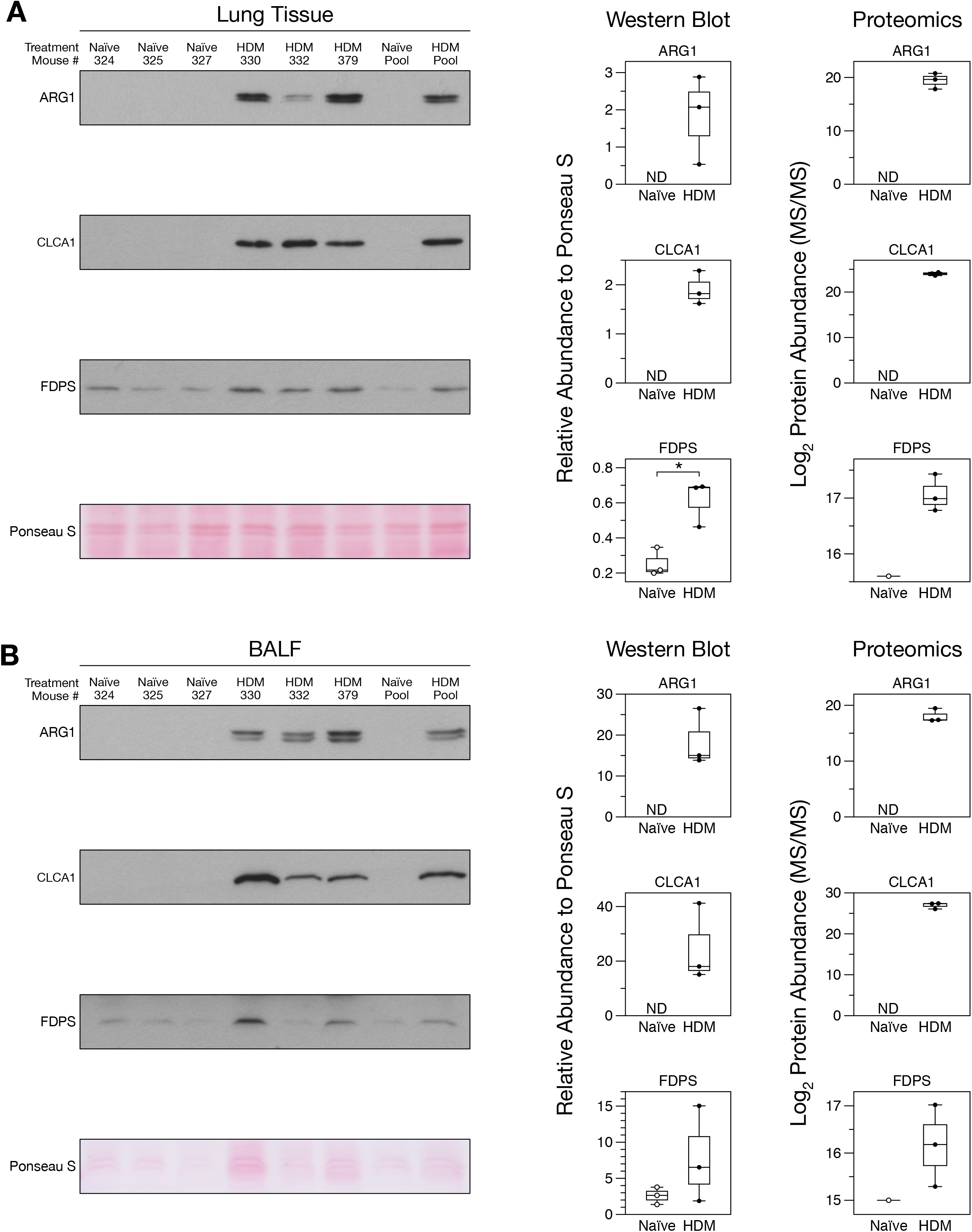
Western blot validation of lung tissue and BALF proteomics. **A-B)** images of scanned chemiluminescent film for ARG1, CLC1, FDPS. Protein bands were normalized to total protein loading (Ponseau S) for quantification. Abbreviations used: Bronchial Alveolar Lavage Fluid (BALF), * (Welch’s unpaired t-test p.adj ≤ 0.05), ND (Not Detected). Protein Names: ARG1 (Arginase-1), CLCA1 (Calcium-activated chloride channel regulator 1), FDPS (Farnesyl pyrophosphate synthase).

### Combining Tissue-BALF proteomes to uncover whole lung responses to HDM challenge

To enable investigation of how HDM-challenge affects protein interactions within the whole lung, we combined all protein IDs from BALF and tissue proteomes (n=2,675). To validate and reduce variability due to missing values, we filtered the dataset to include only those proteins that were detected in all biological replicates for each sample type and experimental condition. This strict criterion reduced the data set to 1,237 proteins for network and pathways analyses. The combined dataset of all tissue and BALF proteins included allergen-naïve (n=558) and HDM-challenged (n=1,064) proteins both containing 385 proteins in common. Using this full dataset we performed PCA to examine the effects of HDM-challenge, and this showed that only 8.1% of the variation in protein abundance correlated with differences between naïve and HDM-treated mice (Y-axis, Figure 3A), whereas 83.8% of variation could be attributed to differences in protein abundance between lung tissue and BALF (X-axis, Figure 3A).

**Figure 3:**
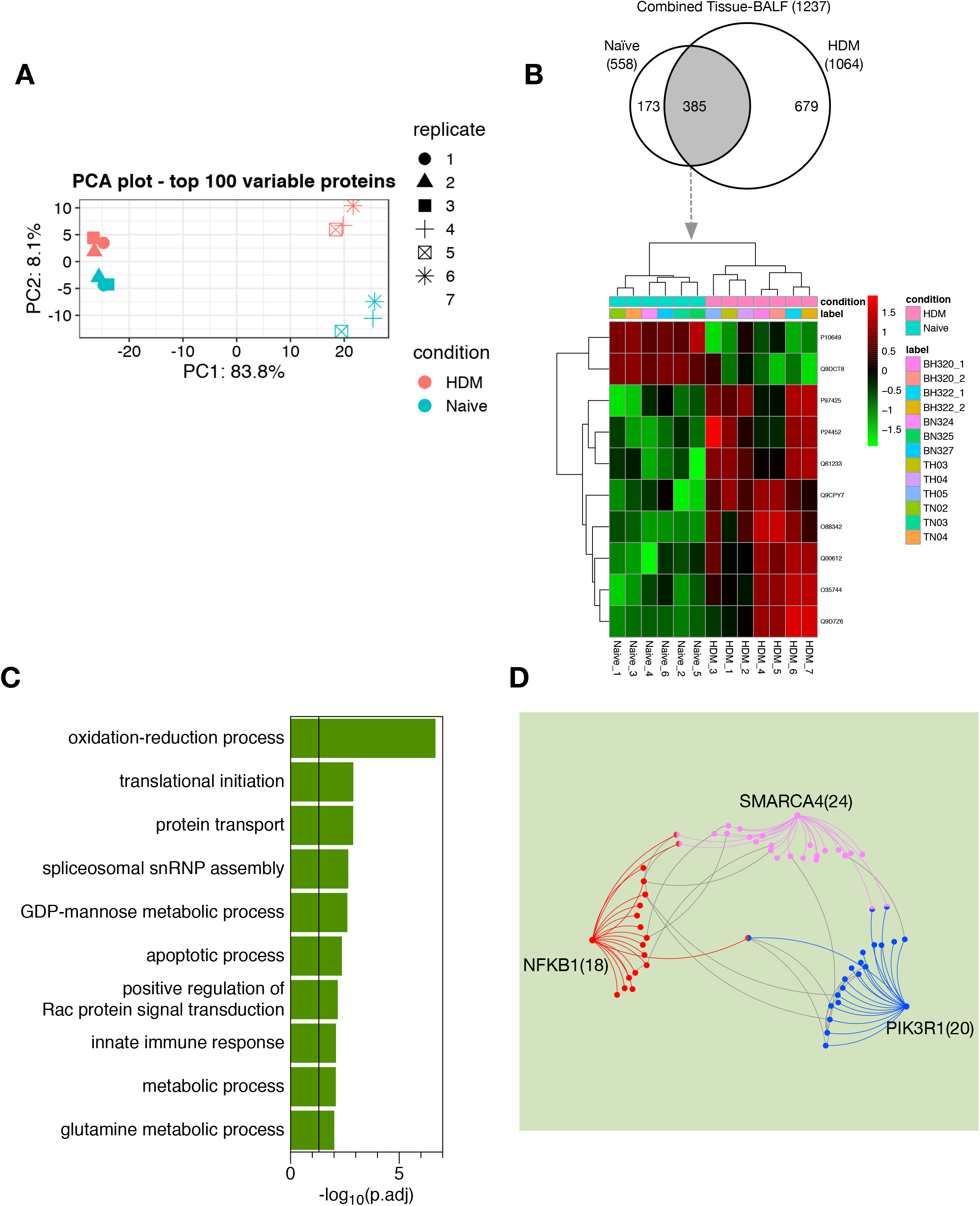
Combining tissue and BALF proteomes identified global processes in the lung of HDM challenged animals. **A)** PCA analysis highlights tissue sample and allergen driven differences in combined proteomes. **B)** Venn diagram showing proteins common to naïve and HDM proteomes and heatmap identifying differentially abundant proteins common to both proteomes. Red – high abundance, Green – low abundance. **C)** Gene ontology biological processes enriched in the HDM proteome (p.adj ≤ 0.05). Vertical line indicates significance threshold. **D)** Protein-protein interaction network of the top 3 most interconnected protein-protein interactions nodes in the HDM proteome. Numbers beside each protein indicate number of direct protein-protein interactions (1^st^ order). Abbreviations used: Bronchial Alveolar Lavage Fluid (BALF). Protein Names: SMARCA4 (Transcription activator BRG1), PIK3R1 (Phosphatidylinositol 3-kinase regulatory subunit alpha), NFKB1 (Nuclear factor NF-kappa-B p105 subunit).

Venn diagram mapping of protein IDs from the combined total lung tissue and BALF dataset showed that 69% of proteins were unique to naïve (173) or HDM-exposed samples (679), and 31% (385) were common to the treatment groups (Figure 3B). Differential expression analysis of the 385 common proteins using LIMMA showed that two proteins were depleted, and eight proteins were enriched after HDM challenge (Table 1 and Figure 3B). To identify which biological processes were represented by these changes, we performed Gene Ontology analysis on a dataset that included the group of eight proteins *enriched by HDM* and the group of 679 proteins that were unique to samples from HDM-challenged mice (see Figure 3B). We identified 36 significantly enriched pathways, with the top 5 biological processes being: “oxidation-reduction process”, “translational initiation”, “protein transport”, “spliceosomal snRNP assembly” and “GDP-mannose metabolic process” (Figure 3C). To complement this analysis, we performed a protein interaction analysis using NetworkAnalyst. The top three nodes, based on the number of connections with other proteins, included: SMARCA4 (Transcription activator BRG1), PIK3R1 (Phosphatidylinositol 3-kinase regulatory subunit alpha) and NFKB1 (Nuclear factor NF-kappa-B p105 subunit) (Figure 3D). This set of networks was most significantly associated with “Interleukin-3, 5 and GM-CSF signaling” (p.adj = 1.40 x10^-8^), forming a backbone for the whole lung response to HDM-challenge.

**Table 1:** Significantly enriched or depleted proteins after HDM challenge in the combined tissue-BALF dataset. Statistical threshold (log_2_FC ≥ 1, p.adj ≤ 0.05). Abbreviations: House Dust Mite (HDM), log_2_ fold change (LogFC)

| Enriched/Depleted in HDM | Uniprot ID | Name | Description | Log<sub>2</sub>FC | p.adj |
| --- | --- | --- | --- | --- | --- |
| Enriched | O35744 | CHIL3 | Chitinase-like protein 3 | 6.89 | 0.0018 |
| Enriched | P97425 | EAR2 | Eosinophil cationic protein 2 | 2.69 | 0.0125 |
| Enriched | Q9D7Z6 | CLCA1 | Calcium-activated chloride channel regulator 1 | 2.68 | 0.0133 |
| Enriched | Q9CPY7 | LAP3 | Cytosol aminopeptidase | 2.44 | 0.00181 |
| Enriched | P24452 | CAPG | Macrophage-capping protein | 2.39 | 0.0172 |
| Enriched | Q61233 | LCP1 | Plastin-2 | 2.08 | 0.00692 |
| Enriched | Q00612 | G6PDX | Glucose-6-phosphate 1-dehydrogenase X | 1.81 | 0.0102 |
| Enriched | O88342 | WDR1 | WD repeat-containing protein 1 | 1.26 | 0.014 |
| Depleted | Q9DCT8 | CRIP2 | Cysteine-rich protein 2 | 2.33 | 0.00181 |
| Depleted | P10649 | GSTM1 | Glutathione S-transferase Mu 1 | 1.59 | 0.00679 |

### Biological significance of proteome changes induced specifically in lung tissue by allergen challenge

To assess lung tissue-specific effects of HDM-challenge we analyzed 1,787 proteins that were unique to all lung tissue samples (Figure 4A). A large fraction of these proteins (n=1,304, 75%) was common to allergen-naïve and HDM-challenged mice, and differential abundance analysis uncovered 43 that were significantly changed after HDM exposure (28 depleted, 17 enriched) (Figure 4B). The proteins most significantly *depleted* by HDM challenge included SERPINA3K (Serine protease inhibitor A3K), CHAD (Chondroadherin) and ADSS (Adenylosuccinate synthetase isozyme 2). We combined the 28 significantly depleted proteins with 138 proteins that were unique to allergen-naïve lung tissue (therefore absent after HDM challenge) (Figure 4A). From this set of proteins that are *uniquely depleted* in the lung by HDM challenge, Gene Ontology analysis revealed that “negative regulation of endopeptidase activity” was the most significantly enriched biological process (Figure 4C).

**Figure 4:**
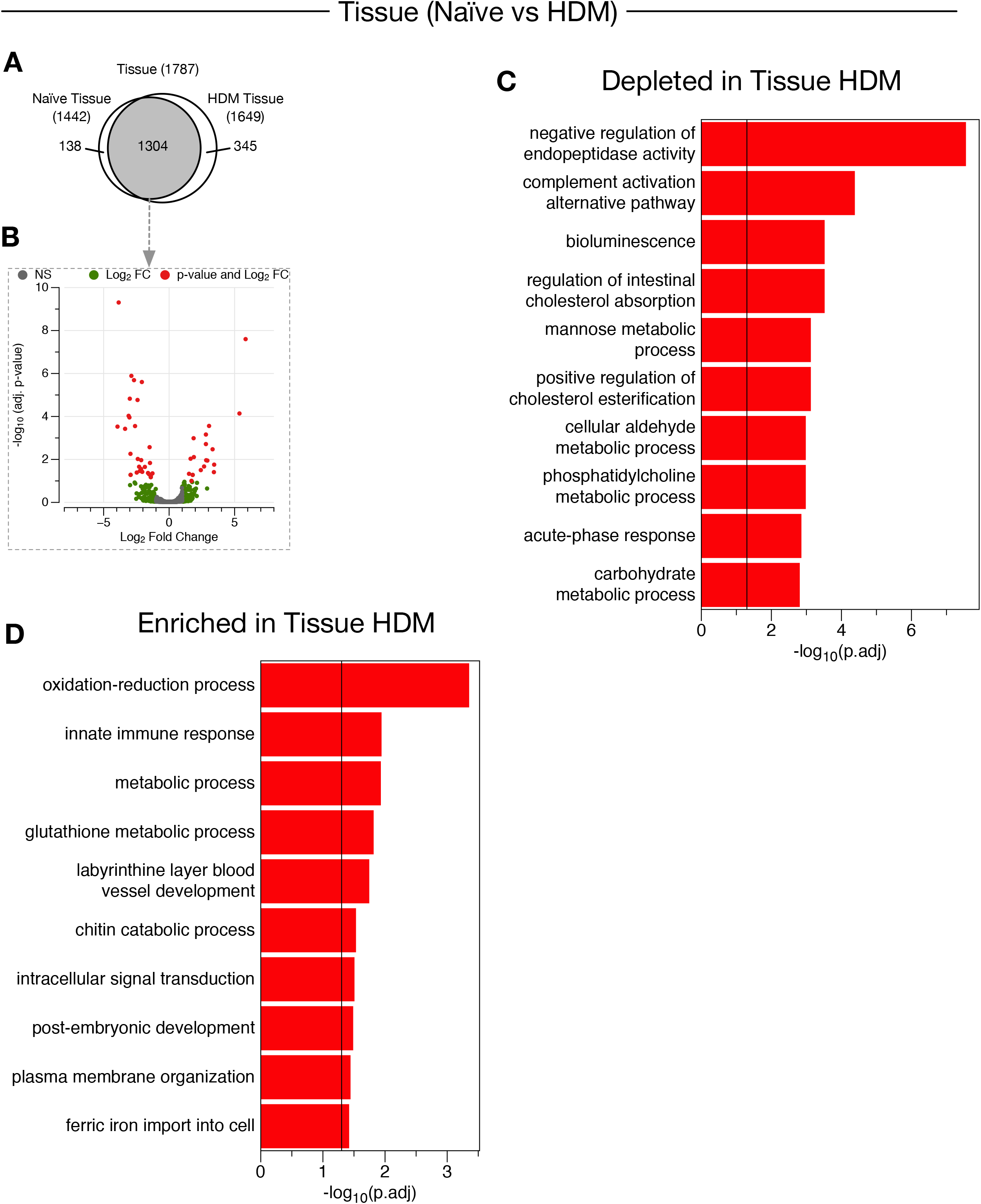
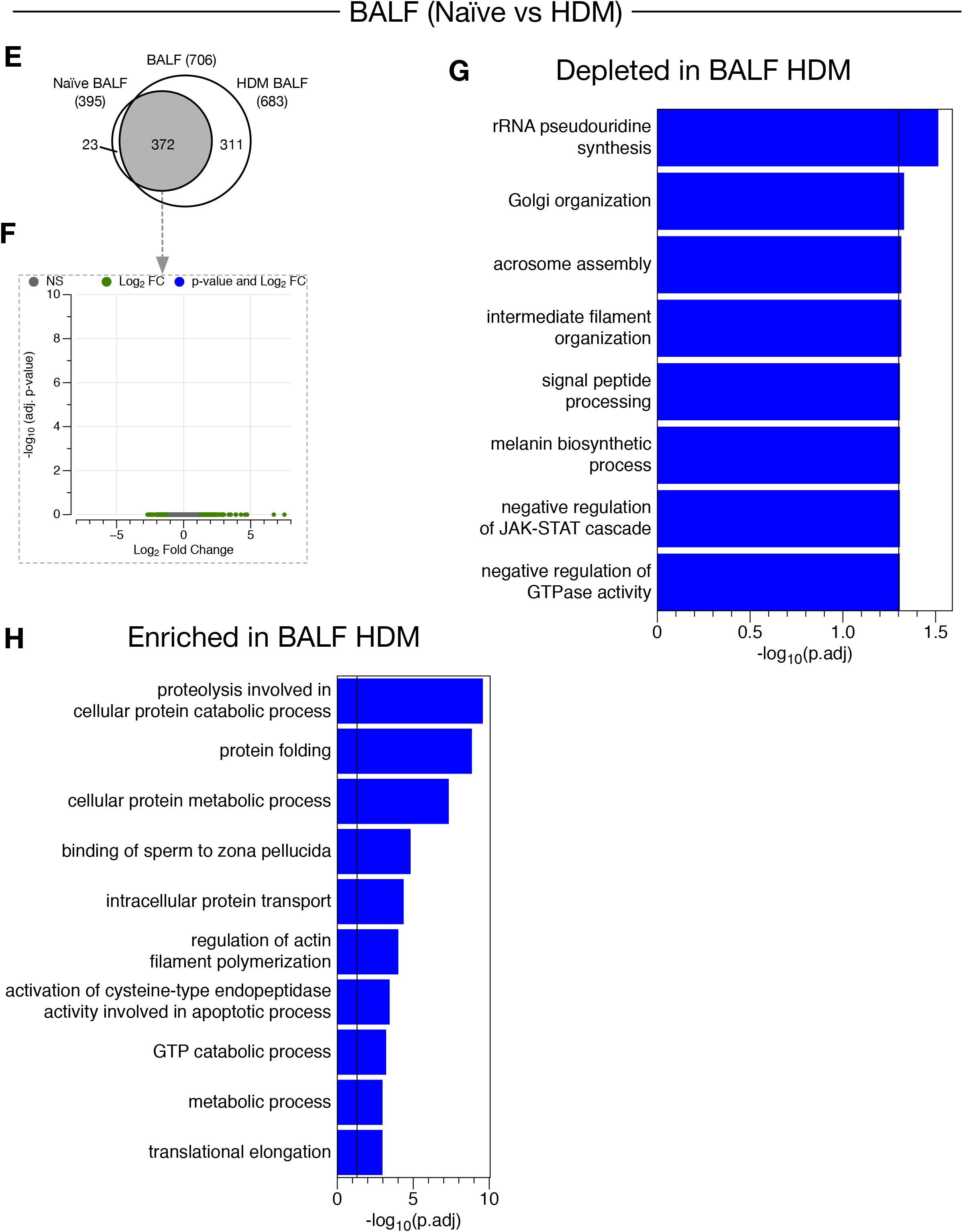
Individually analyzed HDM influenced lung tissue or BALF proteomes do not recapitulate the biology of the Integrated-Tissue-BALF proteome. **A)** Venn diagram of the distribution of protein IDs between allergen-naïve and HDM exposed lung tissue datasets. **B**) Volcano plot showing significantly altered proteins that were common between naïve and HDM tissue proteomes. Red points are significant (p.adj <0.05) and have a log_2_FC > 1. Green points have a log_2_FC > 1 but are not significant. **C-D)** Gene ontology analysis for enriched biological processes on differentially abundant and unique to HDM proteins in tissue. **E)** Venn diagram of the distribution of protein IDs between allergen-naïve and HDM exposed BALF datasets. **F)** Volcano plot showing significantly altered proteins that were common between naïve and HDM BALF proteomes. No significantly altered proteins were detected. **G-H)** Gene ontology analysis for enriched biological processes on unique to HDM proteins in BALF. For brevity, only the Top 10 non-redundant biological processes are shown. Vertical black line indicates significance (p = 0.05).

Of the 17 proteins that were *enriched* (>log_2_ Fold change) by HDM challenge and that were common to allergen-naïve lung tissue samples, CHIL3 (Chitinase-like protein 3), EPX (Eosinophil peroxidase) and LGALS3 (Galectin-3) increased most significantly. We combined the 17 significantly enriched proteins with 345 proteins (Figure 4A) that were unique to HDM-challenged lung tissue samples. Using this set of 362 proteins *uniquely enriched* in lung tissue from HDM-challenged mice, Gene Ontology analysis revealed that “oxidation-reduction” regulation was the most significantly enriched process (Figure 4D).

To extend understanding of the differential abundance of the 362 proteins that we identified as being uniquely altered in allergen exposed lung tissue, we used NetworkAnalyst to develop interactome maps that reveal interaction nodes and signaling networks for biological responses (Figure 5A). The top three interaction nodes for lung tissue were HDAC1 (Histone deacetylase 1), MAPK1 (Mitogen-activated protein kinase 1) and PLCG2 (1-phosphatidylinositol 4,5-bisphosphate phosphodiesterase gamma-2), and these nodes generated an interactome involving multiple pathways. Using the Reactome database linked within NetworkAnalyst, the top 5 pathways were: antigen mediated B-cell receptor activation and secondary messenger generation; hemostasis; signaling by interleukins; proteins associated with G0 and Early G1; and, activation of circadian expression through BMAL1:CLOCK/NPAS2.

**Figure 5:**
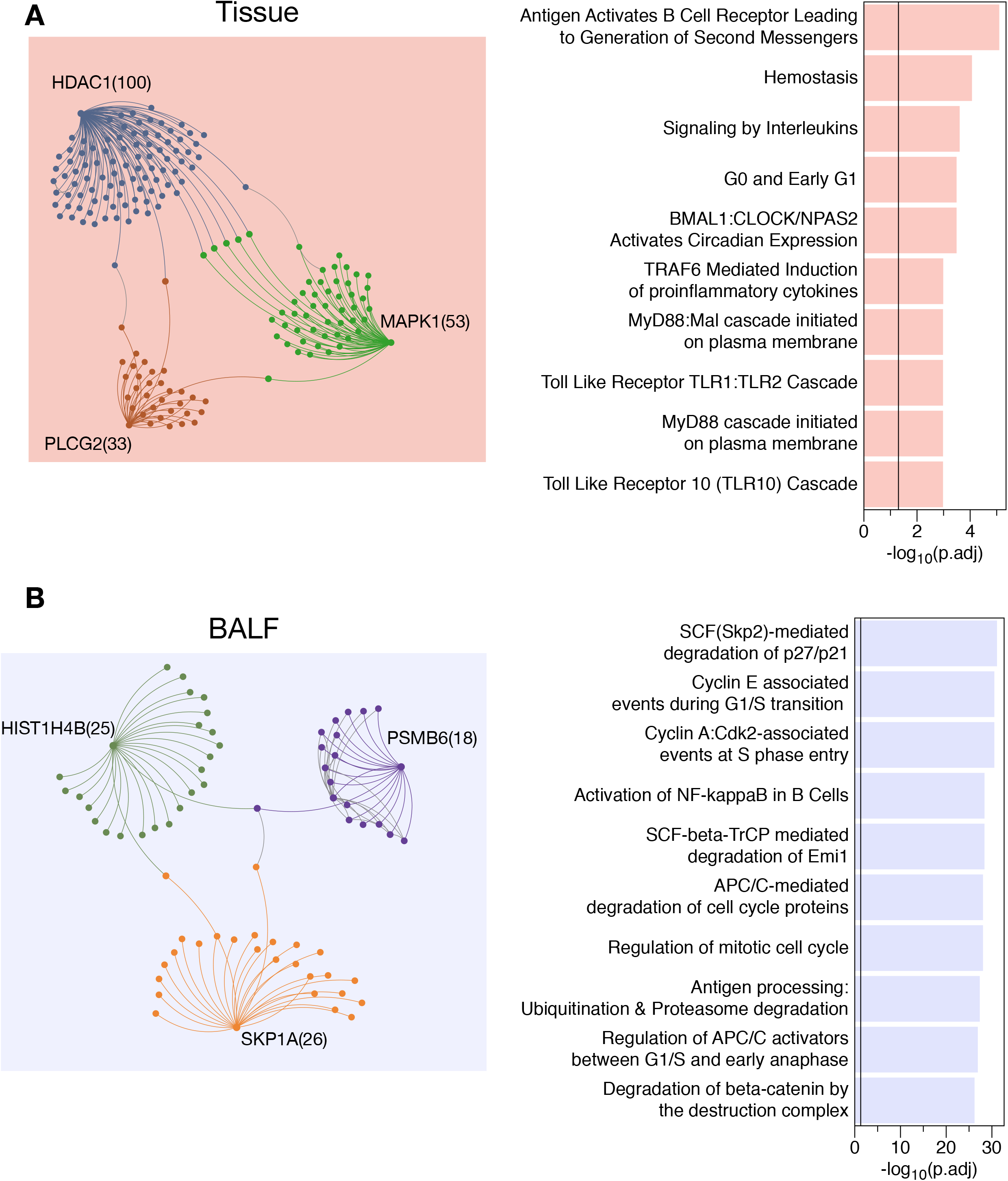
Most connected proteins in the HDM influenced lung tissue and BALF have distinct biological signatures. **A)** Protein-protein interaction network of the top 3 most connected proteins in the HDM influenced lung tissue proteome. **B)** Protein-protein interaction network of the top 3 most connected proteins in the HDM influenced BALF proteome. Biological pathway assessment from each network (Reactome) is shown the right hand side. Protein nodes and lines are coloured to identify direct protein-protein interactions. Other parameters such as node size and line distance is used for illustrative purposes only. Numbers beside each protein indicate number of direct protein-protein interactions (1^st^ order). Abbreviations used: House Dust Mite (HDM), Bronchial Alveolar Lavage Fluid (BALF). Protein Names: HDAC1 (Histone deacetylase 1), MAPK1 (Mitogen-activated protein kinase 1), PLCG2 (1-phosphatidylinositol 4,5-bisphosphate phosphodiesterase gamma-2), SKP1A (S-phase kinase-associated protein 1A), HIST1H4B (H4 clustered histone 2), PSMB6 (Proteasome subunit beta type-6).

### Biological significance of proteome changes induced specifically in BALF by allergen challenge

To assess effects of HDM challenge on the BALF-specific proteome, we analyzed 706 proteins that were unique to all BALF samples (Figure 4E). Approximately half (n=372; 53%) of these protein IDs were common to BALF from allergen-naïve and HDM-challenged mice. Further analysis for differential abundance in these common proteins did not reveal any proteins that were significantly enriched or depleted by allergen challenge (Figure 4F). Therefore, we performed Reactome pathway analysis (linked within NetworkAnalyst) using the 23 proteins that had only been identified in allergen-naïve BALF samples (Figure 4E). This discriminated 10 biological processes that are uniquely *depleted* by HDM challenge in BALF, with “rRNA pseudouridine synthesis” being the most significantly affected (Figure 4G). To determine pathways in BALF that were *enriched* by HDM challenge, we also performed Gene Otology analysis in InnateDB using in the 311 proteins that were unique to BALF from HDM-challenged mice (Figure 4E). This revealed a top 10 biological processes *uniquely enriched* by challenge to inhaled allergen in BALF, and this included “proteolysis involved in cellular protein catabolic process(es)” as the most significantly enriched (Figure 4H).

To extend understanding of the 311 proteins that are uniquely altered in the allergen exposed BALF, we developed interactome maps to identify primary interaction nodes and signaling networks (Figure 5B). Using the Reactome database in NetworkAnalyst, the top three protein-protein interaction (PPI) nodes were SKP1A (S-phase kinase-associated protein 1A), HIST1H4B (H4 clustered histone 2), and PSMB6 (Proteasome subunit beta type-6). This PPI network included diverse pathways, with the top 5 pathways being the SCF(Skp2)-mediated degradation of p27/p21, Cyclin E associated events during G1/S transition, Cyclin A:Cdk2-associated events at S phase entry, activation of NF-κB in B cells, and SCF-beta-TrCP mediated degradation of Emi1.

### Predicting interactions between lung tissue and BALF

To broaden the scope of our study, we interrogated protein interactions across biological compartments using the lung tissue and BALF samples collected from each mouse. To uncover points of integration of biological activities in lung tissue and the surrounding extracellular space in response to allergen challenge, we integrated the 362 proteins that were uniquely enriched by HDM challenge in lung tissue (see Figures 4A, 4B and 5A) with the 311 proteins that were uniquely enriched in BALF (see Figures 4E, 4F and 5B). Importantly, as this integration directly compares the interactions between the lung tissue and BALF, we excluded the 49 proteins that are shared between the two datasets, resulting in 311 and 262 proteins found exclusively in the allergen-exposed lung tissue and BALF datasets respectively. We next identified protein interaction nodes in the combined dataset using NetworkAnalyst, enhancing the rigour of our analyses by filtering the results to only include protein nodes with five or more first-order interactions. From the 37 proteins that met this criterion (12 from lung tissue, 25 from BALF), we selected the top 5 proteins from both the lung tissue and BALF datasets that become enriched in the combined dataset. We found that the integration of BALF proteins with lung tissue-specific proteins resulted in a significant enrichment, 142.1 ± 19.1% (range of 122 - 185%), of first order interactions for nodes that emerged from lung-derived seed proteins (the proteins that start the growth of a network) (Table 2). In contrast, combining lung tissue proteins with those from BALF did not affect the number of interactions for seed protein nodes that originated from BALF (Table 3).

**Table 2:** Network complexity increases when combining the lung tissue and BALF datasets that are uniquely altered after HDM exposure. The number of protein interactions (both 1^st^ and 2^nd^ order) between the lung tissue and BALF datasets are shown (far right column).

| Dataset<br>Uniquely<br>Altered<br>by HDM | Uniprot ID | Name | Description | # of Interactions |  | Protein Node<br>Enrichment in the<br>Combined (%) | Number of 1<sup>st</sup> and 2<sup>nd</sup><br>Order Tissue-BALF<br>Interactions |
| --- | --- | --- | --- | --- | --- | --- | --- |
|  |  |  |  | Tissue | Combined |  |  |
| Tissue | P83917 | CBX1 | Chromobox protein homolog 1 | 4 | 7 | 175.00% | 3 |
| Tissue | P61202 | COPS2 | COP9 signalosome complex<br>subunit 2 | 3 | 5 | 166.67% | 0 |
| Tissue | Q8CBW3 | ABI1 | Abl interactor 1 | 5 | 8 | 160.00% | 0 |
| Tissue | O09106 | HDAC1 | Histone deacetylase 1 | 23 | 34 | 147.83% | 7 |
| Tissue | P70429 | EVL | Ena/VASP-like protein | 5 | 7 | 140.00% | 3 |

To uncover points of crosstalk between lung tissue and BALF proteins we built a combined interaction network using NetworkAnalyst (Figure 6). Once we removed the proteins that are common between the two datasets, we calculated the number of interactions each protein has from within lung tissue (n=311) and BALF (n=262) datasets. We then extracted the top five proteins from the lung tissue that increased as a result of combining with the BALF dataset (Table 2). These included HDAC1, CBX1 (Chromobox protein homolog 1), EVL (Ena/VASP-like protein), ABI1 (Abl interactor 1) and COPS2 (COP9 signalosome complex subunit 2). Since combining these datasets did not enrich the number of interactions from each protein node derived from BALF, for interactome mapping we included the top five seed proteins that we maintained interactions when combined with lung tissue proteins. These included, SPTBN1 (Spectrin beta chain, non-erythrocytic 1), RPS3 (40S ribosomal protein S3), HNRNPAB (Heterogeneous nuclear ribonucleoprotein A/B), PSMA5 (Proteasome subunit alpha type-5), and PSMA2 (Proteasome subunit alpha type-2) (Table 3).

**Figure 6:**
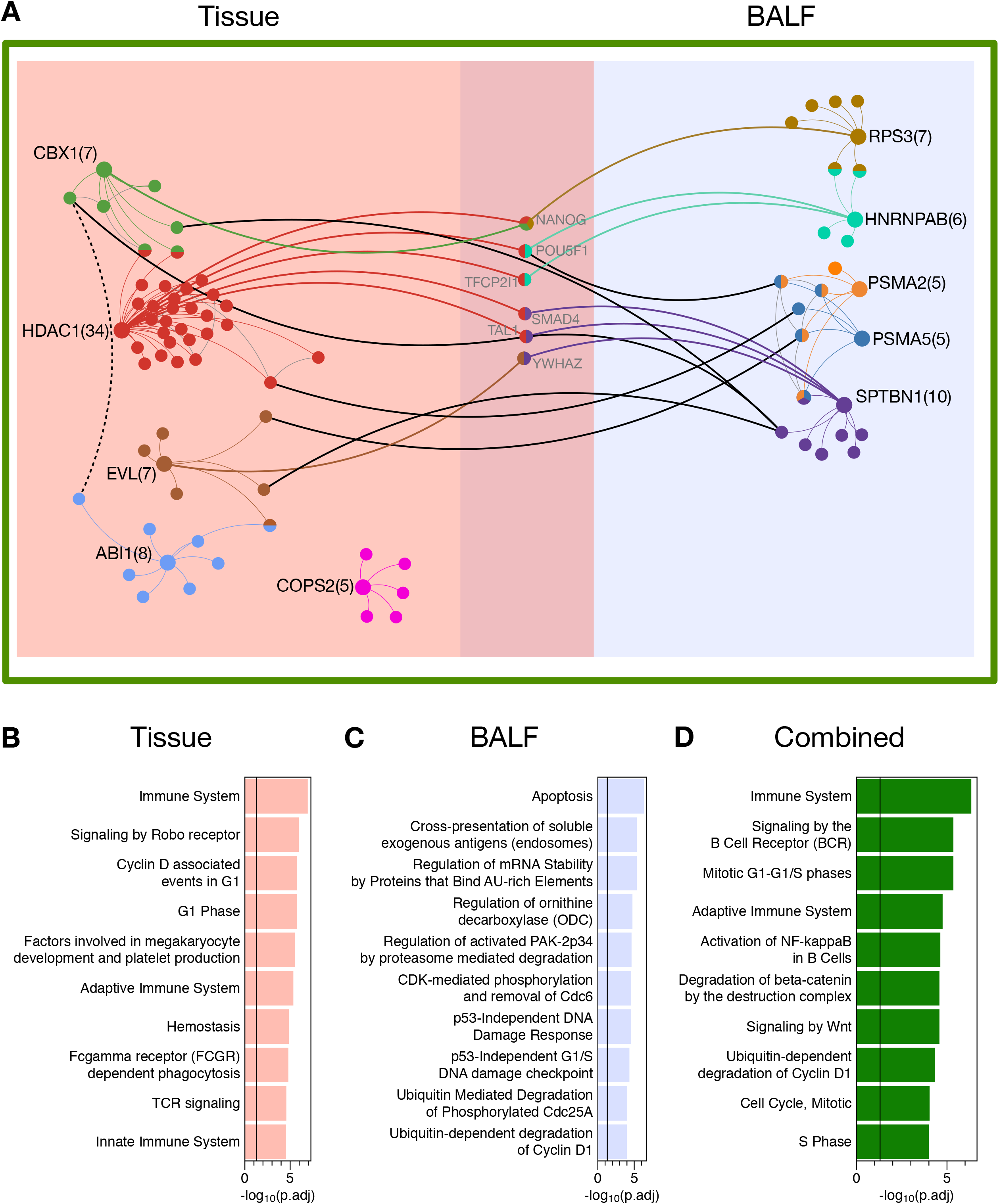
Integrating proteomes not only enrich protein-protein interactions but also connects otherwise separate proteomes together. A) Protein-protein interaction network of the top 5 most connected proteins in the HDM influenced lung tissue and BALF proteomes. Lung tissue nodes (light red) are shown on the left. BALF nodes (light blue) are shown on the right. Protein nodes that are directly involved (1^st^ order) in lung tissue-BALF crosstalk are shown in the center shaded region (purple) and are labelled in grey. The Combined dataset is representative of all protein nodes within green box. B-D) Biological processes derived from the Reactome dataset are colour co-ordinated to their respective networks and each includes the 6 proteins in common (purple). Circles represent individual proteins while protein-protein interactions are shown as lines. Line colour that matches seed nodes indicate direct protein interactions with seed nodes. Circles with split colours indicate shared 1^st^ order interactions between multiple seed nodes. Black lines indicate 2^nd^ order (indirect) inter-dataset lung tissue-BALF crosstalk protein-interactions. Broken lines indicate intra-dataset protein node interactions (2^nd^ order). Number of direct protein-protein interactions (1^st^ order) for each seed node is shown in parentheses. Other parameters such as node size, node distribution and line distance are used for illustrative purposes only. Abbreviations used: Bronchial Alveolar Lavage Fluid (BALF). Protein Names: HDAC1 (Histone deacetylase 1), CBX1 (Chromobox protein homolog 1), EVL (Ena/VASP-like protein), ABI1 (Abl interactor 1), COPS2 (COP9 signalosome complex subunit 2), SPTBN1 (Spectrin beta chain, non-erythrocytic 1), RPS3 (40S ribosomal protein S3), HNRNPAB (Heterogeneous nuclear ribonucleoprotein A/B), PSMA5 (Proteasome subunit alpha type-5), PSMA2 (Proteasome subunit alpha type-2), NANOG (Homeobox protein NANOG), POU5F1 (POU domain, class 5, transcription factor 1), TFCP2I1 (Transcription factor CP2-like protein 1), SMAD4 (Mothers against decapentaplegic homolog 4), TAL1 (T-cell acute lymphocytic leukemia protein 1 homolog) and YWHAZ (14-3-3 protein zeta/delta).

**Table 3:** Network complexity does not change when combining the BALF and lung tissue datasets that are uniquely altered after HDM exposure. The number of protein interactions (both 1^st^ and 2^nd^ order) between the BALF and lung tissue datasets are shown (far right column).

| Dataset<br>Uniquely<br>Altered<br>by HDM | Uniprot ID | Name | Description | # of Interactions |  | Protein Node<br>Enrichment in the<br>Combined (%) | Number of 1<sup>st</sup><br>and 2<sup>nd</sup> Order<br>BALF-Tissue<br>Interactions |
| --- | --- | --- | --- | --- | --- | --- | --- |
|  |  |  |  | BALF | Combined |  |  |
| BALF | Q62261 | SPTBN1 | Spectrin beta chain,<br>non-erythrocytic 1 | 10 | 10 | 100.00% | 7 |
| BALF | P62908 | RPS3 | 40S ribosomal protein<br>S3 | 7 | 7 | 100.00% | 1 |
| BALF | Q99020 | HNRNPAB | Heterogeneous<br>nuclear<br>ribonucleoprotein A/B | 6 | 6 | 100.00% | 2 |
| BALF | Q9Z2U1 | PSMA5 | Proteasome subunit<br>alpha type-5 | 5 | 5 | 100.00% | 2 |
| BALF | Q9R1P3 | PSMB2 | Proteasome subunit<br>beta type-2 | 5 | 5 | 100.00% | 3 |

A unique aspect of our analytical approach is that it uncovered six predicted integration proteins between key pathway nodes in lung tissue and BALF. As shown in Figure 6, these included NANOG (Homeobox protein NANOG), POU5F1 (POU domain, class 5, transcription factor 1), TFCP2I1(Transcription factor CP2-like protein 1), SMAD4 (Mothers against decapentaplegic homolog 4), TAL1 (T-cell acute lymphocytic leukemia protein 1 homolog) and YWHAZ (14-3-3 protein zeta/delta), and each protein had two to four interactions spanning lung tissue and BALF. Among lung-derived seed proteins, HDAC1 interacted with seven predicted integrating proteins, and thus associated with all of the BALF-origin protein nodes. Neither COPS2 nor ABI1 signalling nodes from lung tissue were associated with any of the predicted integration proteins from the BALF dataset, although indirect linkage of ABI1 via EVL and CBX1 is apparent. Among BALF-derived seed nodes, SPTBN1 and HNRNPAB exhibited six and two links, respectively, to predicted integration proteins, thus creating interactions to HDAC1 and EVL in lung tissue. As well, PSMA2 and PSMA5 from BALF projected three interactions to lung tissue nodes (HDAC1 and EVL). Similarly, SPTBN1 in BALF predicted direct linkages to SMAD4, TAL1 and YWHAZ along with indirect linkages between EVL, HDAC1 and CBX1 (Figure 6A). Together, these results reveal that combining proteins in a manner that allows identification of the specific biological compartment from which they were derived enables the critical discrimination of pathways that link the allergic response in lung tissue with that in the surrounding extracellular space.

The impact of combining data for proteins that are uniquely enriched by HDM challenge in matched BALF and lung tissue samples is also evident in the divergence of predicted biological processes that emerges, compared to those based on lung tissue or BALF datasets alone. Figures 6B and 6C list the top 10-biological processes predicted using the Reactome database in NetworkAnalyst for lung tissue and BALF, using the top five seed proteins derived from each, but as influenced by the combined dataset. Prominent lung tissue processes are associated with immune system regulation (innate and adaptive), including T-cell signalling and hemostasis, as well as tissue remodelling (ROBO receptors), and cell division. BALF seed proteins are linked to processes associated with cell stasis (apoptosis, protein turnover, and DNA damage control pathways, including ornithine decarboxylase), as well as antigen presentation. Figure 6D represents the predicted biological processes using all ten seed protein nodes from BALF and lung tissue combined. The list of biological processes reflects a strong dominance by lung tissue-linked processes, in particular immune responses, and the regulation of mitosis and cell division. Of note, two areas that are not evident in lung tissue and BALF alone, but are known components of allergic asthma pathobiology were revealed by the integration of datasets, including B-cell receptor signaling and inflammation, featuring NF-κB activation, pathways induced by Wnt, and degradation of β-catenin.

## Discussion

Murine models of allergic airways inflammation employ inhaled aeroallergen such as HDM to support preclinical and discovery research. Asthma pathobiology involves interaction between recruited inflammatory cells and lung structural cells. In this study we used label-free proteomics and multivariate bioinformatics to describe and compare the molecular interactome in lung tissue and BALF from HDM challenged mice. We used matched samples from individual mice for this process, and in so doing have been able to discriminate responses in tissue and extracellular spaces. Using *in silico* re-integration of datasets, we predict the interactions between the lung tissue and the BALF. We demonstrate that the proteomes of lung tissue and BALF are remarkably different, and this is further enhanced by disparate effects resulting after HDM challenge, highlighting inherent biological diversity in these samples. Examining protein-protein interactions between lung tissue and BALF reveals points for crosstalk between lung tissue and extracellular proteins after HDM exposure. Our study reveals the scope and limits of biological insight that can be obtained from lung tissue or BALF sample proteins alone, and offers potential to interrogate network interactions between the lung tissue and extracellular airway space during allergen challenge.

### Confirming a B cell signaling immune signature in BALF and lung tissue

In the current study, we confirmed that HDM challenge led to predominant eosinophil and neutrophil infiltration of BALF, and we found B cell receptor signaling to be a key component of, and crosstalk between, lung tissue and BALF in allergen exposed mice (Figures 5A and 6D). The immune signature evident in lung tissue and BALF after HDM challenge is predominantly of a Th2-polarizied phenotype, in part dependent on endotoxin levels in HDM^7, 8^. Murine B cells are antigen presenting cells for HDM antigens, and B cell depletion profoundly reduces allergic responses in the lung due to their interactions with resident lung cells and inflammatory cell that infiltrate the lung and airways^9^. Importantly, memory B-cell populations remain in the lung tissue and epithelial cell layers well after HDM sensitization, forming a localized population that can quickly become activated by further allergen challenge^10^. This pattern of a localized memory B cell population contributes to inflammatory responses that associate with airway remodeling in asthmatic patients^11^. As a biological validation of our proteomic investigation, our chief findings reveal a B-cell signaling signature after repeated allergen challenge. Moreover, our network analysis of integrated lung tissue and BALF (collected post-lavage) proteomes reveals multiple levels for crosstalk between the lung tissue and the extracellular microenvironment. In addition to the immunoblot validation of proteins in lung tissue and BALF that we provide, these observations provide evidence of the reliability of our proteome measurement and analysis protocols, and further support an important role for B cells in coordination of biological responses in lung tissue and BALF.

### Protein nodes that link lung tissue and BALF

In lung tissue samples we detected markedly increased HDAC1 (Histone deacetylase 1) after allergen exposure, and it was a seed protein for a major lung tissue interaction node. Moreover, via all six proteins we predicted to integrate lung tissue and BALF responses (Figure 6), HDAC1 appears to interact with all of the BALF protein nodes we identified in HDM-challenge mice. This suggests a fundamental role for HDAC 1 in allergic pathophysiology, which is consistent with its primary role, in conjunction with histone acetyltransferase (HAT), to modulate chromatin accessibility for transcription. Our finding is consistent with evidence from ovalbumin sensitized and challenged mice, in which HDAC1 abundance and activity is increased^12^. Skewed HDAC and HAT activity has been documented in both asthmatic and COPD patients^13, 14^, and HDAC1 activity negatively correlates with disease severity and IL-8 expression^14^. This link with inflammation is due, in part, to transcriptional control of NF-κB signaling and its transcriptional activation of pro-inflammatory genes and antimicrobial peptides^15–17^. Together our study reveals the complexity of HDAC regulation and suggests that modification of transcriptional events leads to physiological and inflammatory changes that span beyond a single biological compartment.

CBX1 abundance was increased in lung tissue after allergen exposure, but exhibited limited interactions with BALF proteins. CBX1 (Chromobox protein homolog 1), a heterochromatin binding protein associated with transcriptional regulation of cell proliferation, also interacts with nuclear lamin (LMNB1) and EMSY (BRCA2-interacting transcriptional repressor)^18, 19^. EMSY, is an interferon-α/β gene regulator^20^, and a genome-wide association study recently identified EMSY (*C11orf30*-LRRC32) to be significantly associated with moderate-to-severe asthma^21^. Similar to CBX1, ABI1 increased significantly in lung tissue after HDM challenge, but network analysis did not reveal interactions with protein nodes in BALF. ABI1 associates with N-WASP to regulate filamentous actin dynamics, as well as acetylcholine mediated contraction of airway smooth muscle^22^. ABI1 is regulated by the tyrosine kinase ABL^22^, which is increased in lungs of asthmatic donors and allergen challenged in mice^23^. Moreover, conditional ABL knockout mice are refractory to allergen challenge-associated increased airway resistance, airway smooth muscle mass, and tracheal contractility^23^. Collectively, our finding of lung tissue-specific induction of CBX1 and ABI1 is consistent with the presence of inherent biological pathways that are primary drivers of fibro-proliferative changes associated with airway remodeling^3^, and airway hyper-responsiveness.

In BALF samples from allergen-challenged mice, SPTBN1 (Spectrin β chain, non-erythrocytic 1; also known as ELF) was increased in abundance, and our network analyses revealed linkage to HDAC1 in tissue via SMAD4 and TAL1. Spectrin is associated with cytoskeletal maintenance with ankyrin and anion exchanger 1 (AE1) to stabilize cortical actin links to the cell membrane^24^. Spectrin repeat and Ph domains of SPTBN1 bind cell membrane phospholipids, thus is positioned to respond to, and effect, interactions between cells in complex biological systems^25, 26^. SPTBN1 also has a critical role in TGF-β induced gene transcription, as SMAD3/4 become mis-localized and dysfunctional in SPTBN1 knockout mouse^27^. This is critical as TGF-β induced SMAD3/4 signaling can induce human airway smooth muscle cell shortening and hyperresponsiveness, and is a primary extracellular driver of asthma pathobiology^28^. Based on its key role in mediating cytoskeletal structure and TGF-β signaling, our finding that SPTBN1 is induced in BALF likely reflects its role in cell-cell interactions, including paracrine signaling systems.

In BALF from allergen exposed mice we detected increased PSMA2 and PSMA5 (proteasome subunit α −2 and −5). These proteins were distinct in that interactions with lung tissue protein nodes were limited to single links with HDAC1 and EVL, suggesting their primary roles lie within the BALF compartment. PSMA2 and PSMA5 maintain structure and function of the 20S proteasomal complex, and has increased activity in BALF from acute respiratory distress patients^29, 30^. Intratracheal instillation of a proteasome inhibitor reduces NF-κB signaling and eosinophil number in the BALF from OVA exposed rats^31^. A definitive role for proteasomal complexes in BALF after allergen challenge is unclear, but they have been implicated in diverse processes such as the formation exogenous peptide antigens for immune cells, degradation of oxidatively damaged proteins, and activation of pre-cursor proteins^32^. We identified PSMA2 and PSMA5 as a significant node in BALF that is relatively disconnected from lung tissue protein nodes, suggesting their principal role lies in regulation of inflammation, involving inflammatory cells, and perhaps their interaction with structural cells of the lung.

### Proteins that mediate lung tissue-BALF crosstalk in allergen exposed mice

We identified remarkable differences in the protein profile between lung tissue and BALF, even more so than the proteome changes that develop in each sample type after allergen challenge. This highlights need to consider a role for protein interactions between lung tissue and BALF after allergen challenge in order to fully interrogate lung system responses to allergic insult. To meet this need, mandated that we use a unique design in which we collected lung tissue and BALF samples from each mouse, performed proteomic analysis in each sample, and then re-combined datasets to predict how the most significant protein nodes in lung tissue and BALF interact. Protein interaction network analysis identified 6 proteins, NANOG, POU5F1, TFCP2I1, SMAD4, TAL1 and YWHAZ, that are present in both lung tissue and BALF datasets, and that could be pivotal mediators of coordinated responses involving both biological compartments.

NANOG (Homeobox protein NANOG) and POU5F1 (POU domain, class 5, transcription factor 1; also known as OCT4) are transcription factors that can function in concert to regulate several genes, including EPHA, FGF2, SMAD1, and SKIL, that are of relevance to lung diseases. These targets regulate cell matrix production and cell motility (EPHA)^33^, induce airway smooth muscle hyperplasia (FGF2)^34^, and provide transcriptional control of TGF-β, either directly by receptor activated SMAD1^35^, and indirectly, by the inhibitory protein, SKIL (also called SnoN)^36^. SKIL, which is part of a family of proteins including SKI and Sno that repress transcription of TGF-β response genes, by interacting with SMAD3 and SMAD4. Interestingly, as noted above, network analysis identified SMAD4, which binds TGF-β induced R-Smads to form transcriptional activator complexes, and TAL1, a hematopoietic transcription factor, as regulators of coordinated responses in BALF and lung tissue. SKI, SKIL and other TGF-β signaling inhibitors regulate cell responses dependent on SMAD4 and TAL1^37, 38^. TGF-β is a central driver of asthma pathobiology, and its elevated levels in lung tissue and BALF is associated with diverse effects. Highlighting this, in mice, though neutralizing antibodies for TGF-β have little effect on HDM challenge-induced lung dysfunction and airway smooth muscle thickening, they do alter the nature and degree of airway inflammation^39^. Overall, our analysis reveals that coordination of responses in BALF and lung tissue compartments involves considerable transcriptional regulation, in particular related to the broad biological effects of TGF-β on tissue repair and remodeling, inflammation, immunity and hematopoietic cell maturation and activation.

Our inter-compartment network analysis also points to the airway epithelium and a key zone for lung tissue-BALF crosstalk, which is consistent with the fact that epithelial barrier function disruption and mucus hypersecretion are hallmark features of the response to repeated allergen inhalation. We identified the transcription factor, TFCP2I1 (Transcription factor CP2-like protein 1; also known as Crtr-1) as a novel co-ordinating protein node. Notably, it has a diverse roles in epithelial cell maturation and modulating β-catenin signaling^40, 41^. To our knowledge, a role for TFCP2I1 in allergic lung pathobiology has not been reported, but in kidneys and salivary glands it is crucial regulator of epithelial cell maturation^40^. There is evidence from lung cancer studies that TFCP2I1 directly interacts with YWHAZ, and protects β-catenin from degradation^42^. YWHAZ, also called 14-3-3ζ, is another protein we predict to be a coordinator of lung tissue-BALF biological responses. It is a binding partner with numerous cytoplasmic and nuclear proteins and modulates signaling associated with apoptosis and cell cycle progression^43, 44^. Furthermore, 14-3-3 proteins can bind the 5’ untranslated region of human surfactant protein A2 mRNA, which may be liked to surfactant deficiency in neonatal respiratory distress syndrome^45^. A role for YWHAZ in β-catenin signaling is potentially important in asthma pathobiology, as β-catenin is associated regulation of TGF-β regulated cellcell adhesion and epithelial to mesenchyme transition^41^, and Wnt/β-catenin signaling is a significant determinant of airway remodeling in asthma^46^. Our finding that TFCP2I1 and YWHAZ may be associated with coincident allergic pathobiology in lung tissue and BALF suggests that the initiation and progression of epithelial barrier disruption and tissue remodeling results from an integrated network of pathways that are associated, in part, with TGF-β and Wnt/β-catenin.

### Limitations

Interpretation of our study is limited by a number of factors. Though we carefully controlled collection methods to reduce variability, such as minimizing lung tissue damage caused by collecting BALF, we cannot discount that BALF samples included a small fraction of cellular proteins. We also only collected samples at one time point (48 hours after final allergen challenge), a strategic choice, as this is when lung dysfunction is greatest. Thus, our work provides only a snapshot of the proteome response, and future studies should adopt a design to assess temporal patterns of the response to HDM challenge. We also only used a single allergen, HDM, but this was a strategic selection as HDM a common human aeroallergen, and has a complex composition that includes multiple allergens, proteases and toxins, therefore offers the advantage of inducing inflammation of a broad profile^7, 8^.

### Summary

We characterized the proteome of lung tissue and BALF from HDM-challenged mice that mimic allergic asthma pathophysiology. Using matched samples from individual animals, our work reveals that lung tissue and BALF proteomes are diverse, and that integrating both datasets reveals additional novel biological processes and protein interaction nodes, in particular highlighting transcriptional regulation as a key integrating parameter for coincident pathobiology in lung tissue and extracellular spaces of the lung. This work provides a resource and approach for identifying new proteins and pathways, and a basis to interrogate interactions between sample compartments to identify mechanisms for airways pathophysiology and, perhaps, new targets for developing therapeutic approaches.

## Methods

### Animal experiments

#### (a) Murine HDM allergen challenge

All animal experiments were planned and performed following the approved protocols and guidelines of the animal ethics board at the University of Manitoba. Female, 8-week old BALB/c mice (6-8 weeks, *n* = 3) were intranasally challenged with HDM (25 μg per mouse, 35 μL saline) five times a week for two weeks (Supplemental Figure S3). HDM extract (Greer Labs; Lenoir, NC) was prepared in sterile phosphate buffered saline (pH 7.4; Life Technologies; Waltham, MA). The HDM extract we used contained 36,000 endotoxin units (EU; 7877 EU/mg of protein or 196.9 EU/dose) and 4.9% Der p 1 protein per vial.

#### (b) Lung function, inflammatory differential cell counts and sample collection

Lung function was performed 48 h after the last HDM challenge. Mice were anesthetized with sodium pentobarbital (90 mg/kg) given intraperitoneally, and tracheotomized with a 20-gauge polyethylene catheter. The catheter was connected to a flexiVent small animal ventilator (Scireq; Montréal, Canada) and mechanically ventilated with a tidal volume of 10 mL/kg body weight, 150 times/min. Forced oscillation technique and positive end expiratory pressure of 3 cmH;O was used for the entire study. A nebulized methacholine (MCh) challenge (0 to 50 mg/mL) was administered to assess concentration dependent response of the respiratory mechanics. Values for each parameter (newtonian resistance, Rn; peripheral tissue damping, G; tissue elastance, H; total resistance, R) were calculated as the peak of all 12 perturbation cycles performed after each MCh challenge. Statistical analysis was conducted using a two-way nested ANOVA with Tukey’s multiple comparison and FDR correction (performed in R).

Post lung function measurement, lungs were lavaged with 1.0 mL of saline two times, for a total of 2 mL containing 0.1% ethylenediaminetetraacetic acid (EDTA; Sigma-Aldrich; St. Louis, MO). BALF was centrifuged to collect the immune cell pellet (1,000 xg, 10 min, 4 °C) and the supernatant was collected and aliquoted prior to flash freezing in liquid nitrogen and storage at −80 °C. Immune cell pellet was resuspended in saline and the total immune cell count was estimated using a hemocytometer. For differential counts, cells were stained with a modified Wright-Giemsa stain (HEMA 3 STAT PACK, Fisher Scientific; Waltham, MA). Cell distribution was analyzed by manually identifying and counting eosinophils, neutrophils, macrophages and lymphocytes in six randomly chosen fields of view examined under a light microscope at 200 x magnification. Post-BAL lung tissue from the left lung and half the right lung were portioned (~35 mg/each), flash frozen in liquid nitrogen and stored at −80 °C.

### Preparation of lung tissue

A randomly selected portioned lung tissue was thawed and weighed before wash the lung tissue of residual blood contamination. Lung tissue was submerged in a 15 mL centrifuge tube containing 15 mL PBS (-CaCl_2_, -MgCl_2_, pH 7.4; Invitrogen, Waltham, MA) with protease/phosphatase inhibitors and mixed in an end-over-end mixer (4 °C, 30 min). Inhibitors including Phenylmethylsulfonyl Fluoride (PMSF, 100 mM stock), Phosphatase Inhibitor Cocktail 2 (Sigma-Aldrich; St. Louis, MO), and Protease Inhibitor (Sigma-Aldrich) each at 1:100 dilution. The tissue surface was cut with scissors to increase effectiveness of the lung tissue wash.

Tissues were homogenized using a TissueRuptor (Qiagen; Venlo, Netherlands) in siliconized centrifuge tubes (Thomas Scientific; Swedesboro, NJ) containing ice cold lysis buffer. Lysis buffer composition: 150 mM NaCl (Fisher Scientific), 50 mM Tris-HCL (pH 7.5; Fisher Scientific, Waltham, MA), 5% glycerol (Sigma-Aldrich), 1% sodium deoxycholate (Sigma-Aldrich), 1% benzonase (25 U/μL; Merck,; Kenilworth, NJ), 1% sodium dodecyl sulfate (SDS; Fisher Scientific), Protease Inhibitor Cocktail 2 (1:100 dilution, Sigma-Aldrich), 1 mM PMSF (Sigma-Aldrich), Phosphatase Inhibitor Cocktail 2 (1:100 dilution, Sigma-Aldrich), 2 mM MgCl_2_ (Fisher Scientific) built up with molecular grade water (Invitrogen). Cell free supernatant (21,000 xg, 10 min, 4 °C, no break) was incubated at room temperature for 30 min to permit benzonase activity before storage at −80°C. All chemicals used for tissue preparation were of molecular/electrophoresis grade.

### Assessment of protein extraction proficiency

From each randomly chosen portion of frozen lung tissue, our protein extraction process yielded an average of 1.146 mg of total protein (μBCA protein assay; Pierce; Waltham, MA). Our rationale for randomizing our selection for a portion of frozen lung tissue resides in the heterogenous nature of allergen induced airway inflammation throughout the lung - there is no region of the lung where the extent of allergen exposure can induce uniform changes across animals. BALF yielded an average of 600 μg of total protein (100 μg per 250 μL aliquot) per mouse (μBCA, Pierce). A coomassie blue stained (GelCode, Invitrogen) gradient LDS-PAGE (NuPAGE; Invitrogen) was imaged (ChemiDock, Bio-Rad) to qualitatively assess total protein molecular weight diversity of both lung tissue (Supplemental Figure S4A) and BALF (Supplemental Figure S4B) protein lysates. No high abundance protein depletion methods were employed to reduce potential elimination bias as shown by the dark band at ~66.5 kDa approximating the molecular weight of albumin (Supplemental Figure S4A,B).

Replicate analysis comparing both technical and biological replicates used mouse BALF HDM sample in parallel protein Filter Assisted Sample Preparation (FASP) procedures. BALF was divided into equal parts (100 μg total protein each) and processed individually and in parallel through our FASP protocol. Our results show that technical variation (Supplemental Figure S4E) is lower than our biological variation (Supplemental Figure S4F) and therefore, we negated the use of technical replicates for our proteomic analysis.

### Filter assisted sample preparation (FASP) of lung tissue and BALF

We modified a previously used FASP protocol for use in lung tissue^47^. Tissue homogenate (300 μg total protein) was supplemented with DTT (final concentration 100 mM) before boiling for 5 min, cooled and supernatant (21,000 xg, 10 min, 4 °C, no break) collected. Tissue homogenates were built with urea buffer (8 M urea, 100 mM Tris) to 800 μL before loading onto 30 kDa molecular weight cut off filters (Amicron Ultra 0.5; Milipore; Burlington, MA). Once samples were loaded (10,000 xg, 10 min, room temperature), columns were washed twice with urea buffer (450 μL; 10,000 xg, 10 min, room temperature). Columns were alkylated (400 μL, 50 mM iodoacidimide in urea buffer; Sigma-Aldrich) for 45 min (protected from light, room temperature) before being stopped by the addition of DTT (20 mM) followed by centrifugation (13,000 xg, 10 min, room temperature). Columns were washed twice (450 μL urea buffer) before a drying the column (14,000 xg, 15 min, room temperature). Trypsin (Trypsin Gold; Promega; Madison, WI) was suspended in digestion buffer (50 mM Tris, 2 mM CaCl_2_, LCMS water) and was added to the column at a protein:trypsin ratio of 50:1. Samples were incubated under shaking conditions at 37 °C for 16 h before being halted by the addition of TFA (1% final concentration) and placing the samples at 4 °C. Columns were incubated (400 μL, 50% methanol, 5 min) prior to being eluted (10,000 xg, 10 min, room temperature). A final wash of the column was also added to the eluted sample (300 μL 15% acetonitrile). Frozen samples (−80 °C) were lyophilized via speed vac and stored at −80 °C.

Mouse BALF samples were processed in a similar manner to that of mouse lung tissue with some minor changes to the protocol. We resuspended 100 μg total protein from thawed BALF supernatant in urea buffer (450 μL; containing 100 mM DTT final concentration) under mixing conditions for 1 h (room temperature) before loading onto columns.

### Peptide desalting by reverse phase 1D-HPLC

Warmed samples were re-suspended in 800 μL TFA (0.5%, LCMS grade water) and centrifuged (21,000 xg, 10 min, 4 °C) to check for undissolved peptides. BALF were loaded onto a C18 column (Luna 10 μM C18(2), 100 Å, 50 x 4.6 mm; Phenomenex, Torrance, CA) while tissue lysate samples were loaded onto a separate C18 column (Phenomenex) offline. Column efficiency and elution conditions were tested using a specialized 6 peptide solution prior to 48 starting the samples^48^. Samples were fractionated across six 1.5 mL tubes and were collected at a flow rate of 500 μL/min with an additional 30 s before and after the eluted peptide spectra was detected. Manual loading was used with no gradient. Samples were frozen at −80 °C. An Agilent 1110 HPLC System using ChemStation for Control and Data Analysis (Santa Clara, CA) was used to analyze the chromatograms from Reverse Phase – HPLC (high pressure liquid chromatography).

### Reconstitution of sample & LC-MS/MS run

Samples were desiccated by speed vac (as previously mentioned) and reconstituted in 50 μL formic acid (0.1%). Peptide concentration was determined by UV spectrophotometry (Nanodrop Spectrophotometer 2000, Thermofisher) at 280 nm. Peptides (2 μg) were diluted in formic acid (0.1%) and injected into an online LC-MS/MS workflow (500 nL/min) using a 3 h gradient run using a Sciex TripleTOF 5600 instrument (Sciex; Framingham, MA). Raw spectra files were converted into Mascot Generic File format (MGF) for protein identification using the tools bundled by the manufacturer. All chemicals used were mass spec grade.

### Data Processing

The MGF files were processed by X!Tandem^49^ against single-missed-cleavage tryptic peptides from the *Mus musclus* Uniprot database (16,704 proteins). The following X!Tandem search parameters were used: 20 ppm and 50 ppm mass tolerance for parent and fragment ions, respectively; constant modification of Cys with iodoacetamide; default set post-translational modifications: oxidation of Met, Trp; N-terminal cyclization at Qln, Cys; N-terminal acetylation, phosphorylation (Ser, Thr, Tyr), deamidation (Asn and Gln); an expectation value cut-off of log_e_ < −1 for both proteins and peptides. Each MS run yielded a list of protein expression values in a log_2_ scale, quantified based on their member peptide MS2 fragment intensity sums. Quantified proteins were detected if at least two non-redundant (unique) peptides with identification scores log_e_ < −1.5 each were identified.

### Data Acquisition: replication, randomization and blocking

In order to homogenize the variance in our studies and avoid confounding technical variability with the biological question, we randomized the order of all our experimental sample work-up. In each treatment group, biological variability was captured with three replicate animals. Moreover, in the case of LC/MS experiments, the sample order was randomized in order to ameliorate effects related to instrumental drift and other variations. The hierarchical order for the magnitude of variability in many omics experiments is well documented and it is this principle that guided our data acquisition strategy^50^.

### Immunoblotting Validation of Proteomics

Protein targets were selected based upon proteomic enrichment after HDM exposure (hypergeometric model with FC >= 2, p.adj <= 0.05, Benjamini-Hochberg multiple comparison) and by antibody availability. Immunoblotting was performed as previously mentioned^51^. Briefly, SDS-PAGE was performed on the same proteins samples used for proteomic analysis. Once 20 μg (total protein) of each sample was transferred onto nitrocellulose membranes, blots were stained with Ponseau S total protein stain (Sigma) prior to blocking at room temperature (5% skim milk, TBST 0.2%, 2 h, RT) and immunoblotted overnight at 4 °C (1% milk, TBST 0.2%). After washing (3 x 15 min, TBST 0.2%), secondary antibody was added (Goat anti-rabbit HRP, 1% milk, TBST 0.2%, 2 h, RT, 1:5000; Sigma). Once washed, blots were flooded with ECL (Amersham; GE Healthcare, Chicago, IL) and bands detected using chemiluminescent film (Hyperfilm ECL; GE Healthcare). Semiquantitative densitometric analysis was performed using scanned film and AlphaEase FC software (Alpha Innotech, San Leandro, CA). Antibodies used: FDPS (PA5-28228, 1:1,000; Thermofisher), ARG1 (ab91279, 1:1,000; Abcam, Cambridge, United Kingdom), CLCA1 (ab180851, 1:2,500; Abcam). To measure average protein content, pooled samples (Naïve Pool and HDM Pool) were prepared by combining equal protein content across biological replicates to equal 20 μg.

### Bioinformatics & Statistical Analysis

Our raw data dataset (n=2,675) across treatment groups: allergen-naïve (n=1,996), and HDM exposed (n=2,502). To limit the number of comparisons we employed a strict filtering criteria; we selected only proteins that are detected in both the compartments, i.e., we kept only those proteins that are identified in all replicates of at least one treatment (HDM or naive). Together this dataset is referred to the Integrated Tissue-BALF dataset (ITB, n=1,237). All the data analyses were conducted in the R, version 3.6.1 (*Action of the Toes*), environment using in-house written code employing packages: limma, vsn, ggplots and pheatmap. The stepwise workflow for the analyses was as follows.

#### (a) Data pre-processing and filtering and normalization

The BALF proteomic measurement of one HDM exposed mouse (labeled #379) was found to be missing over 80% of Uniprot IDs compared to other allergen exposed BALF samples. This sample was also found to be a technical outlier on examination of within group clusters visualized by Principal Component Analysis (PCA). This sample was therefore excluded from all downstream analysis leaving a total of 13 samples to comprise our full dataset (n=13) from which our results are observed (Figure S2B,C). To retain a robust biological signature for the remaining samples, we instituted a data filtering criterion in which we retained proteins whose signal was identified in at least 2 out of 3 technical replicates (or at least 3 out of 4 for HDM BALF) of at least one condition. This then allowed us to impute the rest of the missing values using the k-nearest neighbour (knn) imputation method. To proportionally compare the protein abundances from lung tissue and BALF samples, we performed background correction and normalization by variance stabilizing normalization (vsn)^52^. Using this approach, the variance of the dataset remains nearly constant over the whole dynamic range of our proteomic samples.

#### (b) Exploratory Data Analysis and Differential Analysis (LIMMA)

In order to visualize the (dis)similarities between our proteomics measurements of the samples under different conditions, we used principal component analysis (PCA) as a dimension reduction technique. The variance of all the sample measurements for each protein was calculated, and the top 500 proteins with the greatest variance was selected. We examined the relationship among samples by projecting the original measurements on the two PCs that account for over 80% of the data variance, and plotted these projections as shown in Figure 1A and Figure 3A.

Moreover, in order to identify significantly enriched or depleted proteins we performed differential abundance analysis on the vsn-normalized data using the Linear Models for Microarray Data (LIMMA) approach^53^. We used a statistical threshold of log2 fold changes ≥ 1 and reported FDR adjusted p-values using Benjamini-Hochberg multiple comparison correction. Threshold for statistical significance was p.adj ≤ 0.05. The cutoff criterion for all LIMMA results are uniform across this study.

#### (c) Data visualization

To identify the distribution of individual protein IDs across groups, proportional area venn diagrams were constructed using the eulerr R package (v6.1.0).

Heatmaps were constructed to identify the distribution of protein IDs across conditions. These heatmaps were constructed using either normalized expression values from LIMMA (Figure 1D and 2B) or using a presence/absence classification to identify unique protein IDs (Figure 1B). In addition, grouping analysis (k-means clustering) was constructed for each heatmap identifying groupings across both protein ID and sample condition. Figures were generated using the pheatmap package in R (v1.0.12). Histograms, bargraphs, boxplots, line plots, and linear regressions were constructed using DataGraph (DataGraph v4.5, Visual Data Tools, Inc., Chapel Hill, NC, USA, https://www.visualdatatools.com/).

#### (d) Functional Analyses

Using the Gene Ontology database in InnateDB we first performed over-representation analysis using the default parameters: hypergeometric model with Benjamini-Hochberg multiple comparison correction^54^. To retain non-redundant biological processes we used Revigo with a similarity search pattern of 0.5^55^. Statistical significance of biological pathway enrichment was set at p.adj ≤ 0.05. Moreover, we performed further functional analysis by conducting a protein-protein interaction networks analysis (NetworkAnalyst) and Biological Pathway Assessment (*Reactome*). Protein-protein interaction network analysis was performed by inputting UniprotIDs into NetworkAnalyst (v3.0, accessed April 2020) and analyzed using the InnateDB proteininteraction network database^54, 56^. The top 1^st^ order network (by number of connections) was selected for further analysis. Within each network, the most interconnected protein nodes (from the input Uniprot ID list) that are unique to either BALF or tissue, were selected and subset from the larger networks to examine direct connections between the selected nodes. Biological pathway analysis was then performed using the Reactome database module inside NetworkAnalyst (hypergeometric model with p.adj ≤ 0.05 by Benjamini-Hochberg multiple comparison). The protein UBC not detected in any of our analyses and was removed from all protein-protein interaction analyses.

#### (e) Dataset preparation for predicting lung tissue/BALF interactions

We first integrated the 362 proteins that were uniquely enriched by HDM challenge in lung tissue with the 311 proteins that were uniquely enriched after allergen challenge in BALF. After excluding the 49 proteins that are shared between the two datasets, we input the lung tissue (n=311), BALF (n=262), and combined (n=573) datasets into NetworkAnalyst. To focus on the nodes that are the most interconnected between the BALF and lung tissue datasets we filtered the combined dataset to include proteins with five or more 1^st^ order interactions^57^. From the 37 proteins that met this criterion (12 from lung tissue, 25 from BALF), we selected the top 5 proteins from the BALF and lung tissue dataset that become enriched (% enrichment) in the combined dataset. Selecting the top 5 proteins from each dataset was essential to focus network complexity on the top contributors to protein-interactions.

### Data Storage

We have made all proteomic data freely available in the Mass Spectrometry Interactive Virtual Environment (MassIVE) database housed at UCSD under the id MSV000086003. Files are available at reviewers can grab the files at: ftp://MSV000086003@massive.ucsd.edu (password: a)

## Acknowledgements

Work for this study was supported in part by the CIHR-Canadian Respiratory Research Network (CRRN), Research Manitoba, the DEVOTION Network, and the Biology of Breathing Group, Children’s Hospital Research Institute of Manitoba. THM was supported by studentships from Research Manitoba, CRRN, AllerGen NCE Inc. and Asthma Canada. CDP was supported by a CIHR Banting Fellowship and a Research Manitoba Fellowship. AJ was supported by studentships from Research Manitoba and CRRN. AJH is supported through the Canada Research Chairs Program.

## Author Contributions

THM performed all proteomics experiments, primary and secondary data analysis, prepared all figures and tables and wrote the manuscript draft. CDP provide supervision, guidance and assisted in the completed biostatistics and bioinformatics, design of figures, and editing and writing of the manuscript. TKK provided secondary data quality control, statistical and bioinformatic support as well as support for design and editing of both figures and the final manuscript. AJ and SB were involved with experimental design and completion of animal studies. PE guided sample preparation and performed mass spectrometry. VS contributed to experimental design, primary proteomic data analysis, proteomic data quality control and primary statistics. NM contributed to experimental design and direction, and edited the manuscript. AJH conceived and led design of the study, including scope of biostatistics and bioinformatics, and contributed to writing and editing of the final manuscript.

## Supporting Information

**Supplemental 1:** HDM exposure induces altered lung function and increased inflammatory cell counts. **A)** Lung mechanics were measured 48 h after last HDM treatment using a flexivent small animal ventilator. Increasing doses of methacholine (3-50 mg/mL) intranasally administered to measure changes in airway resistance, tissue resistance and tissue elastance. **B)** Differential immune cell counts from Bronchial Alveolar Lavage Fluid (BALF) of naïve and HDM exposed mice. Each value is representative of the mean and SEM of three biological replicates. Statistical significance was determined by a two-way nested ANOVA with Tukey’s post-hoc test for lung function and unpaired t-test with welch’s correction for cell counts. FDR corrected p-values are reported. *(p.adj ≤ 0.05), **(p.adj ≤ 0.01), ****(p.adj ≤ 0.0001).

**Supplemental 2:** Summary of MS/MS analysis from X!Tandem informatic pipeline **A)** Summary of raw MS/MS data. An EV cutoff of −1.5 and a minimum of two peptides were needed for each peptide and protein ID. **B)** Distribution of unique protein IDs as distributed over the 13 replicates. **C)** Number of protein IDs on a per sample basis. Abbreviations used: Mouse (In House Mouse ID), number of spectra (SPEC), number of peptides (PEPS), number of non-redundant peptides (NR-PEPS), number of proteins (PROTS), number of quantified proteins (QPROT), number of quantified peptides (QPEPS), Mean and Standard Deviation of Log_2_ MS/MS Intensity (MEAN & SD), Bronchial Alveolar Lavage Fluid (BALF), House Dust Mite (HDM). (*) Denotes the biological replicate that was removed due to technical reasons.

**Supplemental 3:** Experimental Workflow. Six mice were split into two groups and exposed to either HDM (House Dust Mite) or PBS (naïve) for two weeks. On day 14, mice were anesthetized and lung function data was collected using a small animal ventilator. Bronchial Alveolar Lavage Fluid (BALF) was collected, spun down to collect immune cells and flash frozen in liquid nitrogen. Post-BAL lung tissue was portioned into 5 equal segments and flash frozen in liquid nitrogen. Lung tissue was washed in PBS to remove excess blood and subsequently homogenized. Samples were processed using FASP, trypsinized, desalted by 1D HPLC and quantified using an online LC-MS/MS proteomic system. Protein IDs were identified using our X!Tandem informatic pipeline and subsequent bioinformatic analysis was conducted on the data. Protein IDs were then filtered retaining ≥ 2/3 or ≥ 3/4 biological replicates per condition prior to variance stabilizing normalization. Protein IDs shared across conditions used normalized protein abundance values as input for differential protein abundance analysis using LIMMA. Protein IDs unique to a condition and those protein IDs which met LIMMA filtering criteria were kept for protein functional analysis using biological pathway enrichment tools (Gene Ontology, Reactome) and protein-protein interaction analysis (NetworkAnalyst). Abbreviations used: Linear Models for Microarray Data (LIMMA), 1D HPLC (1 Dimension High Pressure Liquid Chromatography), PBS (Phosphate Buffered Saline), LC-MS/MS (Liquid Chromatography – Dual Mass Spectrometry), FASP (Filter Assisted Sample Preparation).

**Supplemental 4:** Proteomic quality control and reproducibility tests indicate proteomic variability resides in biological and not technical replicates. **A,B)** Representative SDS-PAGE gels stained for total protein loading with coomassie blue of tissue and BALF protein homogenates (20 μg loading for tissue, 7 μg for BALF). Each lane represents a different mouse. **C,D)** Histograms of detected proteins (mean Log_2_ protein abundance) for the tissue **(C)** and BALF **(D)** datasets are all above the lower limit of detection (vertical black dotted line, 11.67). Technical vs biological replicate assessment of Log_2_ MS/MS intensities from HDM exposed mouse BALF. **E,F)** Biological replicate assessment of Log_2_ MS/MS intensities from mouse BALF. Red line indicates linear goodness of fit (**D,F**).

## References

(1) Calderón, M. A.; Linneberg, A.; Kleine-Tebbe, J.; De Blay, F.; Hernandez Fernandez de Rojas, D.; Virchow, J. C.; Demoly, P. Respiratory allergy caused by house dust mites: What do we really know. J Allergy Clin Immunol. 2015, 136, 38–48.

(2) Choopong, J.; Reamtong, O.; Sookrung, N.; Seesuay, W.; Indrawattana, N.; Sakolvaree, Y.; Chaicumpa, W.; Tungtrongchitr, A. Proteome, Allergenome, and Novel Allergens of House Dust Mite, Dermatophagoides farinae. J Proteome Res. 2016, 15, 422–430.

(3) Jha, A.; Ryu, M. H.; Oo, O.; Bews, H. J.; Carlson, J. C.; Schwartz, J.; Basu, S.; Wong, C. S.; Halayko, A. J. Prophylactic benefits of systemically delivered simvastatin treatment in a house dust mite challenged murine model of allergic asthma. Br J Pharmacol. 2018, 175, 1004–1016.

(4) Wu, J.; Kobayashi, M.; Sousa, E. A.; Liu, W.; Cai, J.; Goldman, S. J.; Dorner, A. J.; Projan, S. J.; Kavuru, M. S.; Qiu, Y.; Thomassen, M. J. Differential proteomic analysis of bronchoalveolar lavage fluid in asthmatics following segmental antigen challenge. Mol Cell Proteomics. 2005, 4, 1251–1264.

(5) O’Neil, S. E.; Sitkauskiene, B.; Babusyte, A.; Krisiukeniene, A.; Stravinskaite-Bieksiene, K.; Sakalauskas, R.; Sihlbom, C.; Ekerljung, L.; Carlsohn, E.; Lötvall, J. Network analysis of quantitative proteomics on asthmatic bronchi: effects of inhaled glucocorticoid treatment. Respir Res. 2011, 12, 124.

(6) Burg, D.; Schofield, J. P. R.; Brandsma, J.; Staykova, D.; Folisi, C.; Bansal, A.; Nicholas, B.; Xian, Y.; Rowe, A.; Corfield, J.; Wilson, S.; Ward, J.; Lutter, R.; Fleming, L.; Shaw, D. E.; Bakke, P. S.; Caruso, M.; Dahlen, S. E.; Fowler, S. J.; Hashimoto, S.; Horváth, I.; Howarth, P.; Krug, N.; Montuschi, P.; Sanak, M.; Sandström, T.; Singer, F.; Sun, K.; Pandis, I.; Auffray, C.; Sousa, A. R.; Adcock, I. M.; Chung, K. F.; Sterk, P. J.; Djukanović, R.; Skipp, P. J.; The, U.-B. S. G. Large-Scale Label-Free Quantitative Mapping of the Sputum Proteome. J Proteome Res. 2018, 17, 2072–2091.

(7) Pascoe, C. D.; Jha, A.; Basu, S.; Mahood, T.; Lee, A.; Hinshaw, S.; Falsafi, R.; Hancock, R. E. W.; Mookherjee, N.; Halayko, A. J. The importance of reporting house dust mite endotoxin abundance: impact on the lung transcriptome. Am J Physiol Lung Cell Mol Physiol. 2020, 318, L1229–L1236.

(8) Piyadasa, H.; Altieri, A.; Basu, S.; Schwartz, J.; Halayko, A. J.; Mookherjee, N. Biosignature for airway inflammation in a house dust mite-challenged murine model of allergic asthma. Biol Open. 2016, 5, 112–121.

(9) Wypych, T. P.; Marzi, R.; Wu, G. F.; Lanzavecchia, A.; Sallusto, F. Role of B cells in T_H_ cell responses in a mouse model of asthma. J Allergy Clin Immunol. 2018, 141, 1395–1410.

(10) Turner, D. L.; Goldklang, M.; Cvetkovski, F.; Paik, D.; Trischler, J.; Barahona, J.; Cao, M.; Dave, R.; Tanna, N.; D’Armiento, J. M.; Farber, D. L. Biased Generation and In Situ Activation of Lung Tissue-Resident Memory CD4 T Cells in the Pathogenesis of Allergic Asthma. J Immunol. 2018, 200, 1561–1569.

(11) Svenningsen, S.; Kirby, M.; Starr, D.; Coxson, H. O.; Paterson, N. A.; McCormack, D. G.; Parraga, G. What are ventilation defects in asthma. Thorax. 2014, 69, 63–71.

(12) Su, X. M.; Ren, Y.; Li, M. L.; Zhao, X.; Kong, L. F.; Kang, J. Performance evaluation of histone deacetylases in lungs of mice exposed to ovalbumin aerosols. J Physiol Pharmacol. 2018, 69, 265–273.

(13) Ito, K.; Caramori, G.; Lim, S.; Oates, T.; Chung, K. F.; Barnes, P. J.; Adcock, I. M. Expression and activity of histone deacetylases in human asthmatic airways. Am J Respir Crit Care Med. 2002, 166, 392–396.

(14) Ito, K.; Ito, M.; Elliott, W. M.; Cosio, B.; Caramori, G.; Kon, O. M.; Barczyk, A.; Hayashi, S.; Adcock, I. M.; Hogg, J. C.; Barnes, P. J. Decreased histone deacetylase activity in chronic obstructive pulmonary disease. N Engl J Med. 2005, 352, 1967–1976.

(15) Poynter, M. E.; Irvin, C. G.; Janssen-Heininger, Y. M. Rapid activation of nuclear factor-kappaB in airway epithelium in a murine model of allergic airway inflammation. Am J Pathol. 2002, 160, 1325–1334.

(16) Kallsen, K.; Andresen, E.; Heine, H. Histone deacetylase (HDAC) 1 controls the expression of beta defensin 1 in human lung epithelial cells. PLoS One. 2012, 7, e50000.

(17) Redhu, N. S.; Saleh, A.; Lee, H. C.; Halayko, A. J.; Ziegler, S. F.; Gounni, A. S. IgE induces transcriptional regulation of thymic stromal lymphopoietin in human airway smooth muscle cells. J Allergy Clin Immunol. 2011, 128, 892–896.e2.

(18) Huang, Y.; Myers, M. P.; Xu, R. M. Crystal structure of the HP1-EMSY complex reveals an unusual mode of HP1 binding. Structure. 2006, 14, 703–712.

(19) Lu, J.; Gilbert, D. M. Proliferation-dependent and cell cycle regulated transcription of mouse pericentric heterochromatin. J Cell Biol. 2007, 179, 411–421.

(20) Ezell, S. A.; Polytarchou, C.; Hatziapostolou, M.; Guo, A.; Sanidas, I.; Bihani, T.; Comb, M. J.; Sourvinos, G.; Tsichlis, P. N. The protein kinase Akt1 regulates the interferon response through phosphorylation of the transcriptional repressor EMSY. Proc Natl Acad Sci U S A. 2012, 109, E613–21.

(21) Chen, J.; Chen, Q.; Wu, C.; Jin, Y. Genetic variants of the C11orf30-LRRC32 region are associated with childhood asthma in the Chinese population. Allergol Immunopathol (Madr). 2019, 48, 390–394.

(22) Wang, T.; Cleary, R. A.; Wang, R.; Tang, D. D. Role of the adapter protein Abi1 in actin-associated signaling and smooth muscle contraction. J Biol Chem. 2013, 288, 20713–20722.

(23) Cleary, R. A.; Wang, R.; Wang, T.; Tang, D. D. Role of Abl in airway hyperresponsiveness and airway remodeling. Respir Res. 2013, 14, 105.

(24) Nestor, M. W.; Cai, X.; Stone, M. R.; Bloch, R. J.; Thompson, S. M. The actin binding domain of ßI-spectrin regulates the morphological and functional dynamics of dendritic spines. PLoS One. 2011, 6, e16197.

(25) Ray, S.; Chakrabarti, A. Membrane interaction of erythroid spectrin: surface-density-dependent high-affinity binding to phosphatidylethanolamine. Mol Membr Biol. 2004, 21, 93100.

(26) Diakowski, W.; Sikorski, A. F. Interaction of brain spectrin (fodrin) with phospholipids. Biochemistry. 1995, 34, 13252–13258.

(27) Tang, Y.; Katuri, V.; Dillner, A.; Mishra, B.; Deng, C. X.; Mishra, L. Disruption of transforming growth factor-beta signaling in ELF beta-spectrin-deficient mice. Science. 2003, 299, 574–577.

(28) Ojiaku, C. A.; Cao, G.; Zhu, W.; Yoo, E. J.; Shumyatcher, M.; Himes, B. E.; An, S. S.; Panettieri, R. A. TGF-β 1 Evokes Human Airway Smooth Muscle Cell Shortening and Hyperresponsiveness via Smad3. Am J Respir Cell Mol Biol. 2018, 58, 575–584.

(29) Keller, I. E.; Vosyka, O.; Takenaka, S.; Kloß, A.; Dahlmann, B.; Willems, L. I.; Verdoes, M.; Overkleeft, H. S.; Marcos, E.; Adnot, S.; Hauck, S. M.; Ruppert, C.; Günther, A.; Herold, S.; Ohno, S.; Adler, H.; Eickelberg, O.; Meiners, S. Regulation of immunoproteasome function in the lung. Sci Rep. 2015, 5, 10230.

(30) Sixt, S. U.; Adamzik, M.; Spyrka, D.; Saul, B.; Hakenbeck, J.; Wohlschlaeger, J.; Costabel, U.; Kloss, A.; Giesebrecht, J.; Dahlmann, B.; Peters, J. Alveolar extracellular 20S proteasome in patients with acute respiratory distress syndrome. Am J Respir Crit Care Med. 2009, 179, 1098–1106.

(31) Elliott, P. J.; Pien, C. S.; McCormack, T. A.; Chapman, I. D.; Adams, J. Proteasome inhibition: A novel mechanism to combat asthma. J Allergy Clin Immunol. 1999, 104, 294–300.

(32) Sixt, S. U.; Peters, J. Extracellular alveolar proteasome: possible role in lung injury and repair. Proc Am Thorac Soc. 2010, 7, 91–96.

(33) Yamazaki, T.; Masuda, J.; Omori, T.; Usui, R.; Akiyama, H.; Maru, Y. EphA1 interacts with integrin-linked kinase and regulates cell morphology and motility. J Cell Sci. 2009, 122, 243–255.

(34) Jeon, S. G.; Lee, C. G.; Oh, M. H.; Chun, E. Y.; Gho, Y. S.; Cho, S. H.; Kim, J. H.; Min, K. U.; Kim, Y. Y.; Kim, Y. K.; Elias, J. A. Recombinant basic fibroblast growth factor inhibits the airway hyperresponsiveness, mucus production, and lung inflammation induced by an allergen challenge. J Allergy Clin Immunol. 2007, 119, 831–837.

(35) Liu, F.; Hata, A.; Baker, J. C.; Doody, J.; Cárcamo, J.; Harland, R. M.; Massagué, J. A human Mad protein acting as a BMP-regulated transcriptional activator. Nature. 1996, 381, 620–623.

(36) Nyman, T.; Trésaugues, L.; Welin, M.; Lehtiö, L.; Flodin, S.; Persson, C.; Johansson, I.; Hammarström, M.; Nordlund, P. The crystal structure of the Dachshund domain of human SnoN reveals flexibility in the putative protein interaction surface. PLoS One. 2010, 5, e12907.

(37) He, J.; Tegen, S. B.; Krawitz, A. R.; Martin, G. S.; Luo, K. The transforming activity of Ski and SnoN is dependent on their ability to repress the activity of Smad proteins. J Biol Chem. 2003, 278, 30540–30547.

(38) Meng, X.; Lu, P.; Bai, H.; Xiao, P.; Fan, Q. Transcriptional regulatory networks in human lung adenocarcinoma. Mol Med Rep. 2012, 6, 961–966.

(39) Fattouh, R.; Midence, N. G.; Arias, K.; Johnson, J. R.; Walker, T. D.; Goncharova, S.; Souza, K. P.; Gregory, R. C.; Lonning, S.; Gauldie, J.; Jordana, M. Transforming growth factor-beta regulates house dust mite-induced allergic airway inflammation but not airway remodeling. Am J Respir Crit Care Med. 2008, 177, 593–603.

(40) Yamaguchi, Y.; Yonemura, S.; Takada, S. Grainyhead-related transcription factor is required for duct maturation in the salivary gland and the kidney of the mouse. Development. 2006, 133, 4737–4748.

(41) Giangreco, A.; Lu, L.; Vickers, C.; Teixeira, V. H.; Groot, K. R.; Butler, C. R.; Ilieva, E. V.; George, P. J.; Nicholson, A. G.; Sage, E. K.; Watt, F. M.; Janes, S. M. ß-Catenin determines upper airway progenitor cell fate and preinvasive squamous lung cancer progression by modulating epithelial-mesenchymal transition. J Pathol. 2012, 226, 575–587.

(42) Chen, C. H.; Chuang, S. M.; Yang, M. F.; Liao, J. W.; Yu, S. L.; Chen, J. J. A novel function of YWHAZ/ß-catenin axis in promoting epithelial-mesenchymal transition and lung cancer metastasis. Mol Cancer Res. 2012, 10, 1319–1331.

(43) Weerasekara, V. K.; Panek, D. J.; Broadbent, D. G.; Mortenson, J. B.; Mathis, A. D.; Logan, G. N.; Prince, J. T.; Thomson, D. M.; Thompson, J. W.; Andersen, J. L. Metabolic-stress-induced rearrangement of the 14-3-3ζ interactome promotes autophagy via a ULK1- and AMPK-regulated 14-3-3ζ interaction with phosphorylated Atg9. Mol Cell Biol. 2014, 34, 4379–4388.

(44) Lee, J. J.; Lee, J. S.; Cui, M. N.; Yun, H. H.; Kim, H. Y.; Lee, S. H.; Lee, J. H. BIS targeting induces cellular senescence through the regulation of 14-3-3 zeta/STAT3/SKP2/p27 in glioblastoma cells. Cell Death Dis. 2014, 5, e1537.

(45) Noutsios, G. T.; Ghattas, P.; Bennett, S.; Floros, J. 14-3-3 isoforms bind directly exon B of the 5’-UTR of human surfactant protein A2 mRNA. Am J Physiol Lung Cell Mol Physiol. 2015, 309, L147–57.

(46) Kumawat, K.; Koopmans, T.; Gosens, R. β-catenin as a regulator and therapeutic target for asthmatic airway remodeling. Expert Opin Ther Targets. 2014, 18, 1023–1034.

(47) Wisniewski, J. R.; Zougman, A.; Nagaraj, N.; Mann, M. Universal sample preparation method for proteome analysis. Nat Methods. 2009, 6, 359–362.

(48) Krokhin, O. V.; Spicer, V. Peptide retention standards and hydrophobicity indexes in reversed-phase high-performance liquid chromatography of peptides. Anal Chem. 2009, 81, 9522–9530.

(49) Craig, R.; Beavis, R. C. TANDEM: matching proteins with tandem mass spectra. Bioinformatics. 2004, 20, 1466–1467.

(50) Karakach, T. K.; Wentzell, P. D.; Walter, J. A. Characterization of the measurement error structure in 1D 1H NMR data for metabolomics studies. Anal Chim Acta. 2009, 636, 163–174.

(51) Tran, T.; McNeill, K. D.; Gerthoffer, W. T.; Unruh, H.; Halayko, A. J. Endogenous laminin is required for human airway smooth muscle cell maturation. Respir Res. 2006, 7, 117.

(52) Välikangas, T.; Suomi, T.; Elo, L. L. A systematic evaluation of normalization methods in quantitative label-free proteomics. Brief Bioinform. 2018, 19, 1–11.

(53) Ritchie, M. E.; Phipson, B.; Wu, D.; Hu, Y.; Law, C. W.; Shi, W.; Smyth, G. K. limma powers differential expression analyses for RNA-sequencing and microarray studies. Nucleic Acids Res. 2015, 43, e47.

(54) Breuer, K.; Foroushani, A. K.; Laird, M. R.; Chen, C.; Sribnaia, A.; Lo, R.; Winsor, G. L.; Hancock, R. E.; Brinkman, F. S.; Lynn, D. J. InnateDB: systems biology of innate immunity and beyond--recent updates and continuing curation. Nucleic Acids Res. 2013, 41, D1228–33.

(55) Supek, F.; Bošnjak, M.; Škunca, N.; Šmuc, T. REVIGO summarizes and visualizes long lists of gene ontology terms. PLoS One. 2011, 6, e21800.

(56) Zhou, G.; Soufan, O.; Ewald, J.; Hancock, R. E. W.; Basu, N.; Xia, J. NetworkAnalyst 3.0: a visual analytics platform for comprehensive gene expression profiling and meta-analysis. Nucleic Acids Res. 2019, 47, W234–W241.

(57) Vallabhajosyula, R. R.; Chakravarti, D.; Lutfeali, S.; Ray, A.; Raval, A. Identifying hubs in protein interaction networks. PLoS One. 2009, 4, e5344.

