## Supplementary figures and images for "Integrating Proteomes for Lung Tissue and Lavage Reveals Pathways that Link Responses in Allergen-Challenged Mice"

### Supplemental S1

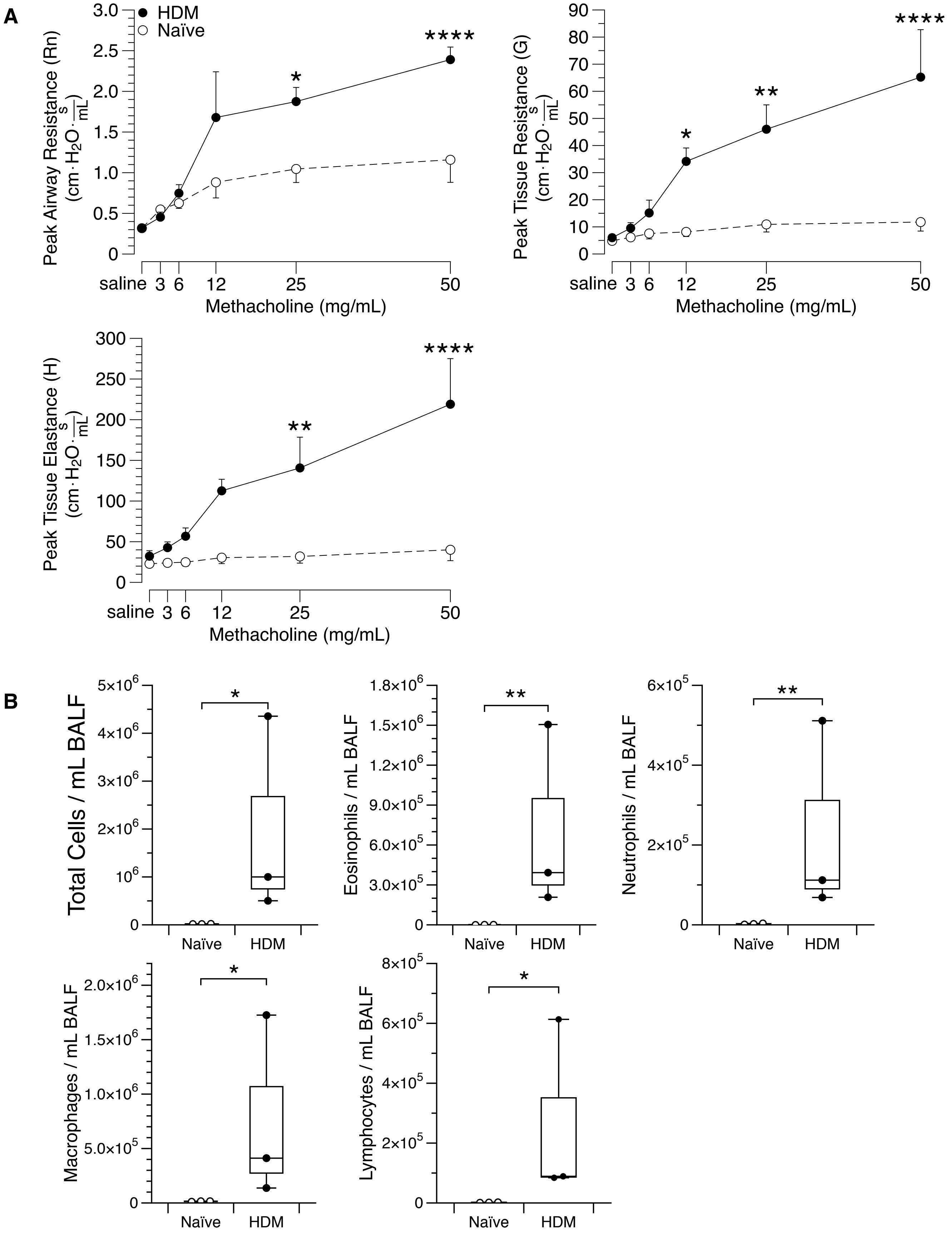

### Supplemental S3

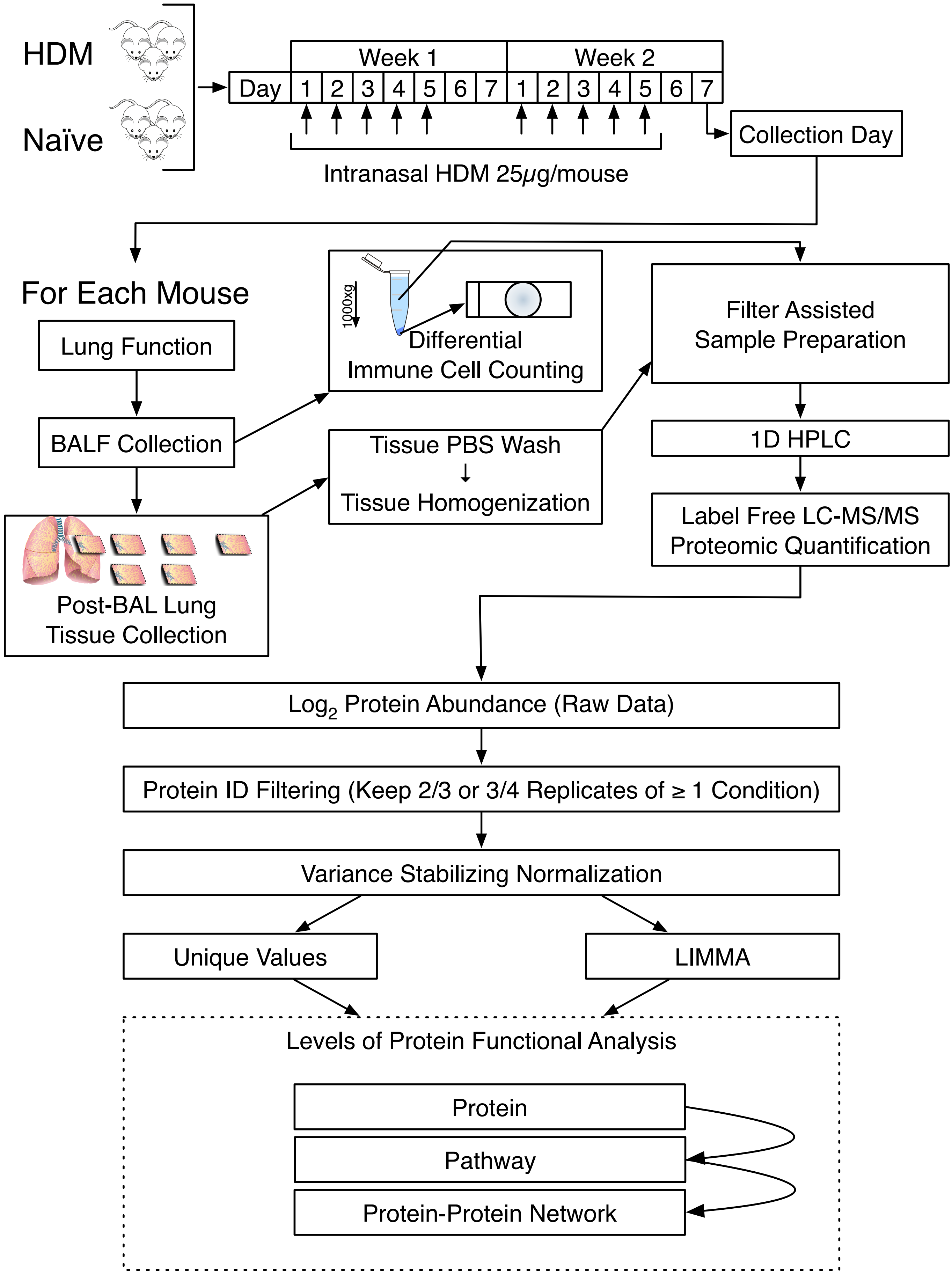

### Supplemental S4

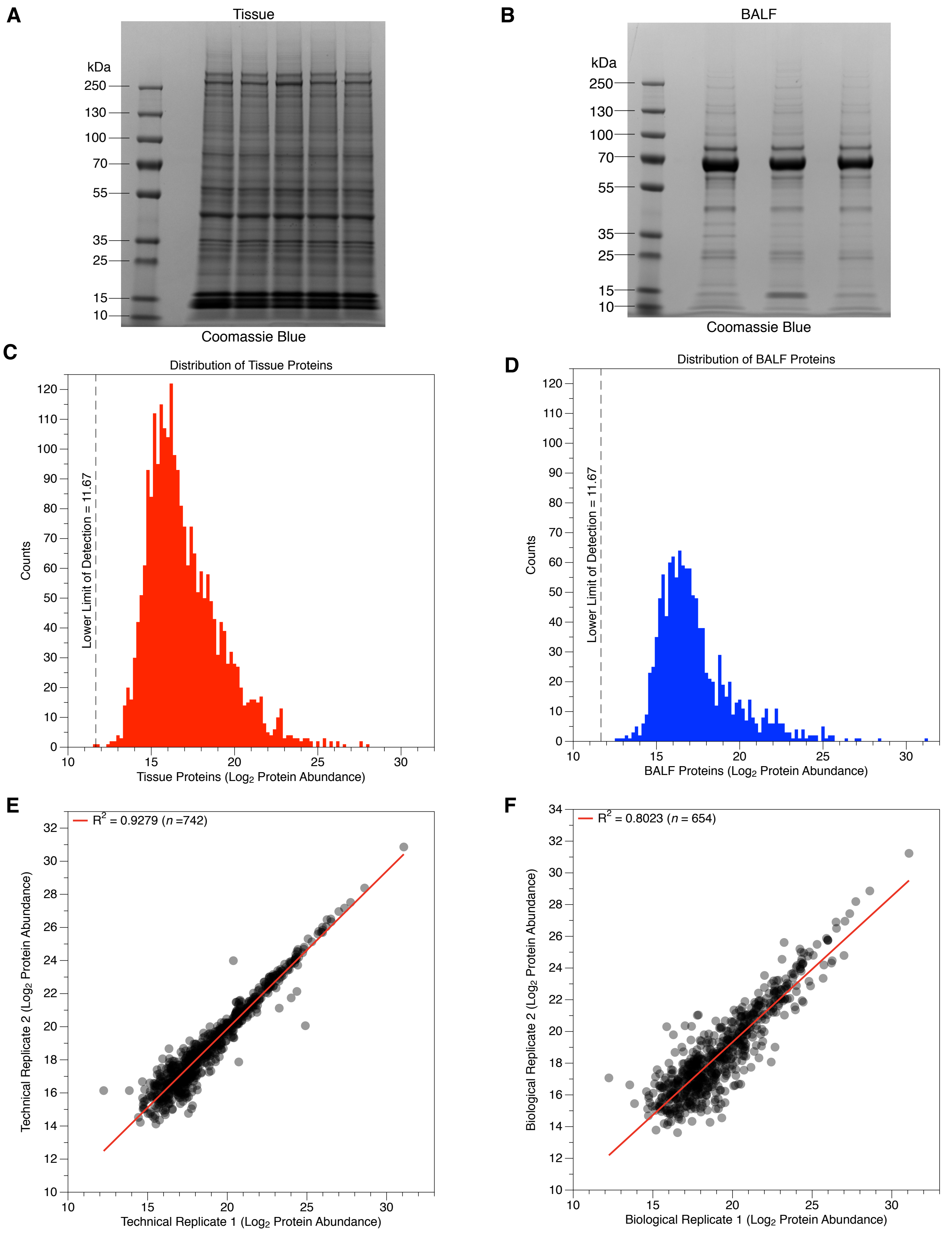
