## Supplemental S2 for "Integrating Proteomes for Lung Tissue and Lavage Reveals Pathways that Link Responses in Allergen-Challenged Mice"

Figure S2

**A**

| | Source | Treatment | Mouse | Replicate # | SPEC | PEPS | NR-PEPS | PROTS | QPROT | QPEPS | MEAN | MIN - MAX |
|---|---|---|---|---|---|---|---|---|---|---|---|---|
| | Tissue | Naïve | 324 | 1 | 64168 | 37718 | 11742 | 2445 | 1463 | 10119 | 17.89 | 12.52 - 28.38 |
| | Tissue | Naïve | 325 | 2 | 63445 | 38682 | 12151 | 2479 | 1521 | 10586 | 17.65 | 12.70 - 28.27 |
| | Tissue | Naïve | 327 | 3 | 62555 | 36558 | 11348 | 2458 | 1438 | 9702 | 17.56 | 12.43 - 28.56 |
| | Tissue | HDM | 330 | 1 | 63362 | 36771 | 12619 | 2676 | 1635 | 10838 | 17.76 | 13.05 - 27.98 |
| | Tissue | HDM | 332 | 2 | 64143 | 37685 | 12819 | 2686 | 1657 | 11113 | 17.79 | 12.67 - 28.11 |
| | Tissue | HDM | 379 | 3 | 67747 | 45155 | 14904 | 2832 | 1852 | 13230 | 17.64 | 11.66 - 27.69 |
| | BALF | Naïve | 324 | 1 | 59853 | 34520 | 4389 | 784 | 444 | 3837 | 18.03 | 13.27 - 31.29 |
| | BALF | Naïve | 325 | 2 | 59183 | 31939 | 3909 | 745 | 394 | 3375 | 18.07 | 12.77 - 31.58 |
| | BALF | Naïve | 327 | 3 | 60175 | 32051 | 3985 | 785 | 434 | 3459 | 18.18 | 12.02 - 31.65 |
| | BALF | HDM | 330 | 1 | 62067 | 30537 | 8186 | 1541 | 897 | 7125 | 18.44 | 12.24 - 31.08 |
| | BALF | HDM | 330 | 2 | 61747 | 25904 | 7796 | 1543 | 876 | 6706 | 18.47 | 13.29 - 30.86 |
| | BALF | HDM | 332 | 3 | 60590 | 28341 | 6972 | 1345 | 783 | 6047 | 18.25 | 13.51 - 31.23 |
| | BALF | HDM | 332 | 4 | 59517 | 27213 | 6687 | 1315 | 735 | 5760 | 18.16 | 12.69 - 30.98 |
| * | BALF | HDM | 379 | 5 | 65701 | 34853 | 5517 | 988 | 563 | 4844 | 17.63 | 11.74 - 30.80 |

**B**

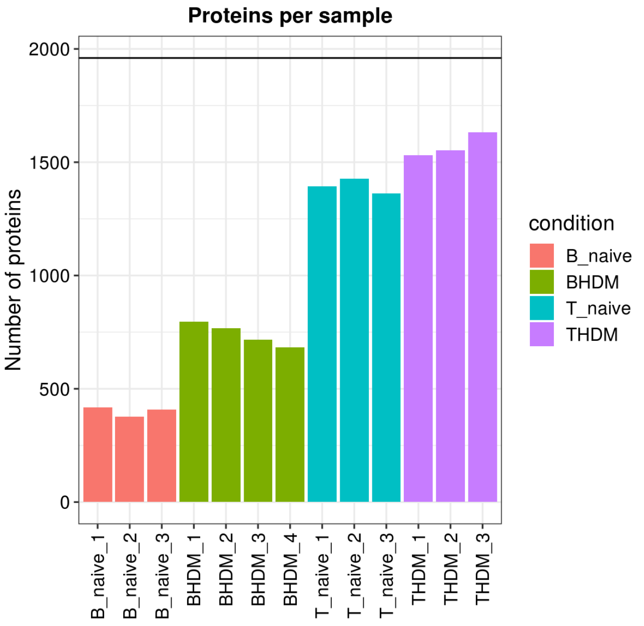


**C**

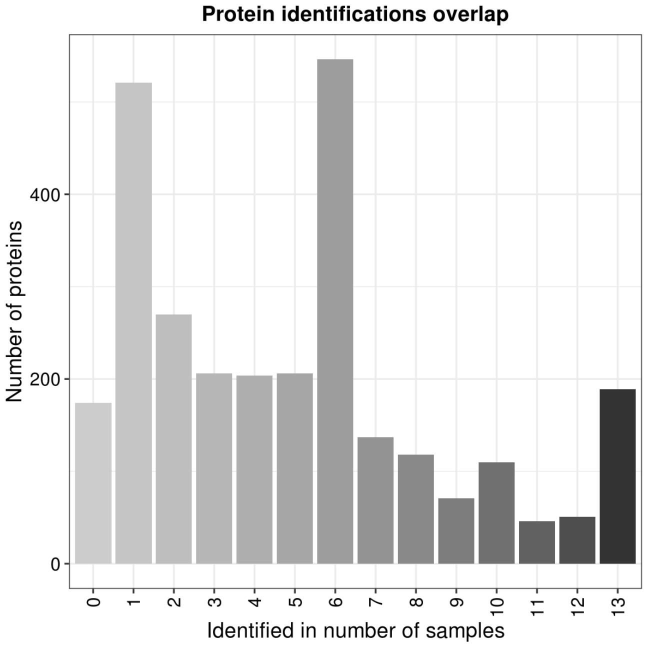
